# Dimethyl fumarate modulates the Duchenne muscular dystrophy disease program following short-term treatment in *mdx* mice

**DOI:** 10.1101/2022.09.15.508124

**Authors:** Cara A. Timpani, Stephanie Kourakis, Danielle A. Debruin, Dean G. Campelj, Nancy Pompeani, Narges Dargahi, Angelo P. Bautista, Ryan M. Bagaric, Elya J. Ritenis, Lauren Sahakian, Patricia Hafner, Peter G. Arthur, Jessica R. Terrill, Vasso Apostolopoulos, Judy B. de Haan, Nuri Guven, Dirk Fischer, Emma Rybalka

## Abstract

New medicines are urgently required to treat the fatal neuromuscular disease, Duchenne muscular dystrophy (DMD). DMD involves progressive muscle damage and weakness, which are preceded by oxidative stress, inflammation, and mitochondrial dysfunction. Dimethyl fumarate (DMF) is a potent small molecule nuclear erythroid 2-related factor 2 (Nrf2) activator with current clinical utility in the treatment of multiple sclerosis and psoriasis. Pharmaceutical targeting of Nrf2 by DMF has strong translational potential for DMD, given it: (1) promotes antioxidant defence systems; (2) has a potent immuno-modulatory profile; and (3) can be rapidly re-purposed into clinical care strategies for DMD patients. Here, we tested two weeks of daily 100mg/kg DMF versus 5mg/kg standard care prednisone (PRED) treatment during the peak muscle degeneration period in juvenile *mdx* mice, the gold standard murine DMD model. Both drugs modulated seed genes driving the DMD disease program and improved muscle force production in fast-twitch muscle. However, only DMF showed pro-mitochondrial effects that protected contracting muscles from fatigue, improved histopathology and augmented clinically compatible muscle function tests. In contrast, PRED treatment stunted mouse growth, worsened histopathology and modulated many normally expressed inflammatory and extracellular matrix (ECM) genes consistent with pan immunosuppression. These findings suggest DMF could be a more selective modulator of the DMD disease program with better efficacy and fewer side effects than standard care PRED therapy warranting follow-up studies to progress clinical translation.

## Introduction

Drug re-purposing is an efficient strategy to deliver medicines to market in a time- and cost-effective manner. Rare diseases could benefit most from this strategy because they are often fatal, rapidly progressive and have high unmet clinical need (1). Duchenne muscular dystrophy (DMD) is a devastating neuromuscular disease that matches these criteria making it a good candidate for drug re-purposing. In DMD, muscles lack functional dystrophin protein from the cytoskeleton due to mutation of the longest human gene, *Dmd*. This deficiency results in muscle fragility, dysregulated ion channels and a complex pathophysiology leading to the chronic degeneration of skeletal muscles (reviewed extensively in (2)). Cardiac and smooth muscle are also affected, as well as other tissue types expressing dystrophin isoforms (e.g., vascular endothelium, brain), although to a lesser extent. DMD patients are wheelchair bound by ~12 years and ultimately die from cardiorespiratory failure in early adulthood (~26 years). Corticosteroids (i.e., prednisone (PRED)/prednisolone, deflazacort) have prevailed as standard care pharmacotherapy for >20 years and are associated with severe side effects making them unsuitable for some patients (3). Emerging therapeutics targeted at the genetic mechanism offer new hope, but robust dystrophin expression leading to clear and consistent functional benefits remains problematic (4). The scope for gene therapies to slow disease course will likely increase as they are prescribed for younger patients who are still ambulatory (5), however, maximum efficacy may depend on multi-modal treatments that also target the underlying pathobiology. There remain few treatment options for advanced DMD patients.

Dimethyl fumarate (DMF) is a small molecule immunomodulator clinically used to treat relapsing remitting multiple sclerosis (RRMS; Tecfidera®) and psoriasis (Fumaderm®). Both diseases are driven by auto-immunity and DMF effectively treats this aetiology. Alternative therapeutic applications are currently under investigation for chronic diseases that share similar aetiology in both clinical and pre-clinical studies (reviewed in (6)). DMF’s established mechanism of action is through activation of the transcription factor, nuclear erythroid factor-2 related factor 2 (Nrf2), which incites the cytoprotective program against toxic stress (summarised in Fig 1). This program results in anti-oxidation, anti-inflammation, and detoxification, and is especially effective in the immune system to control overactivation. Additional complementary mechanisms (shown in Fig 1) include: (1) agonism of the hydroxycarboxylic acid receptor 2 (HCAR2), to inhibit membrane breakdown and resolve inflammation (7); (2) inhibition of the glycolytic enzyme, glyceraldehyde 3-phosphate dehydrogenase (GAPDH), which rewires metabolism through the mitochondria (8); and (3) blocking toll-like receptor 4 induction of pro-inflammatory cytokines (9). Following oral consumption, DMF is rapidly converted in the gut to its bioactive form, monomethyl fumarate, which is taken up by cells and converted to fumarate. Fumarate causes succinylation of key cytosolic proteins, including negative repressor of Nrf2, kelch-like ECH-associated protein 1 (KEAP1), resulting in Nrf2 activation (10). It also stimulates mitochondrial oxidative phosphorylation through integration into the Kreb’s cycle where it is ultimately completely metabolised (11).

**Figure 1.**
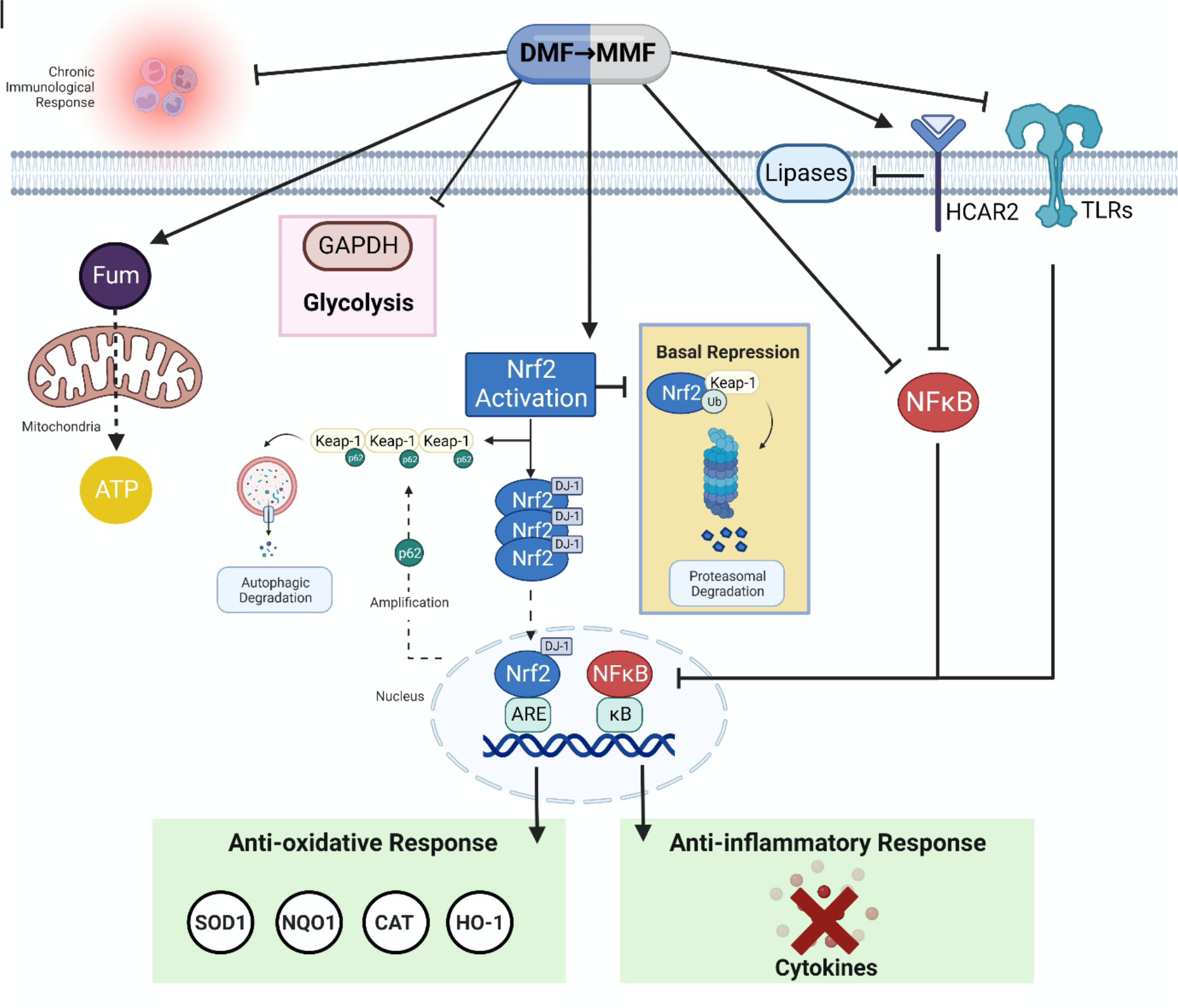
Mechanisms of action of dimethyl fumarate (DMF). DMF is rapidly converted to its bioactive form, monomethyl fumarate (MMF) in the gut and circulated to tissues. Inside cells, MMF is converted to fumarate, which binds Keap1, resulting in dissociation of the Keap1-Nrf2 complex. When complexed in the basal state, Keap1 represses Nrf2 activity by targeting the complex for degradation by the ubiquitin proteosome. Once dissociated from Keap1, DJ-1 chaperones Nrf2 into the nucleus where Nrf2 binds the antioxidant response element (ARE) resulting in transcription of antioxidant genes for superoxide dismutase 1 (SOD1), NAD(P)H dehydrogenase (quinone) 1 (NQO1), catalase (CAT) and haemoxygenase 1 (HO-1). Meanwhile, Keap1 is sequestered by p62, which initiates autophagy and amplifies Nrf2-mediated ARE transcription. Fumarate also inhibits master controller of inflammation, nuclear factor kappa B (NF-κB), which suppresses nuclear binding of κB and transcription of cytokines that drive the inflammatory response. MMF also inhibits NF-κB via agonism of the hydroxycarboxylic acid receptor 2 (HCAR2) and antagonism of toll-like receptors (TLRs) on the membrane. Fumarate causes metabolic shifts by inhibiting glyceraldehyde-3-phosphate dehydrogenase (GAPDH) activity, and therefore, glycolysis. Fumarate enters mitochondria via the malate-aspartate shuttle where it is ultimately sequestered into the matrix tricarboxylic acid cycle where it is completely metabolised to yield ATP and CO_2_.

Investigational therapeutics for DMD have historically fallen short in clinical trials (4) highlighting both the complexity of the pathophysiological milieu that drives degeneration of dystrophin-deficient muscles and disparity between the human condition and animal models. Pre-clinical drug investigations primarily use the gold standard *mdx* mouse, which recapitulates a milder DMD phenotype than human patients (12) resulting in poor translation of experimental treatments into the clinic. Despite inducing robust muscle preservation in pre-clinical *mdx* trials using sexually mature, disease-stable mice, promising drugs have so far been unsuccessful to attenuate disease progression in clinical trials (reviewed in (13) using myostatin inhibitors as an example). Severe damage bouts during the juvenile and senile periods in *mdx* mice could and should be leveraged to assess more human comparable disease. We recently developed a theoretical context for Nrf2’s suitability as a candidate drug target to treat DMD (14). Mitochondrial function, autophagy, satellite cell cycling, calcium homeostasis and inflammation are all chronically dysregulated in DMD and once activated, Nrf2 can positively modulate these processes to promote cell survival. Indeed, knocking Nrf2 out of *mdx* mice escalates DMD pathology when disease is aggravated by running (15). Importantly, applying the Nrf2 activators, sulforaphane (16–18) and curcumin (19) to the *mdx* mouse lessens myopathy. However, no study has investigated an Nrf2 activator drug with clinical indication that could be rapidly translated for the clinical management of DMD nor contrasted drug efficacy against standard care glucocorticoids. In this study, we aimed to pre-clinically evaluate the efficacy of DMF against standard care PRED using juvenile *mdx* mice with severe spontaneous onset MD. Our data demonstrate DMF as a translational candidate for future clinical trials.

## Results

### DMF is well tolerated in mice and improves muscle function test performance but not blood biomarkers of DMD

Because the gold standard *mdx* mouse manifests an overall milder DMD phenotype compared to humans, we used a juvenile period of rapid growth and muscle damage to test the tolerability and effects of short-term DMF treatment compared to standard care PRED (and 0.5% methyl cellulose vehicle (VEH)) using clinically compatible function tests and fluid biomarkers (Fig 2A). Two weeks of DMF treatment had no impact on animal welfare indices, including growth rate, and food and water consumption (Supp Fig 1A-C) or on body weight corrected organ mass (data not shown). In contrast, PRED stunted growth from 12-14 days (Supp Fig 1A), increased water consumption between 10-11 days (Supp Fig 1C) and reduced spleen mass (data not shown). Our data are consistent with the known growth inhibiting, mineralocorticoid, and immunosuppressive side-effects of PRED treatment in children (20).

**Figure 2:**
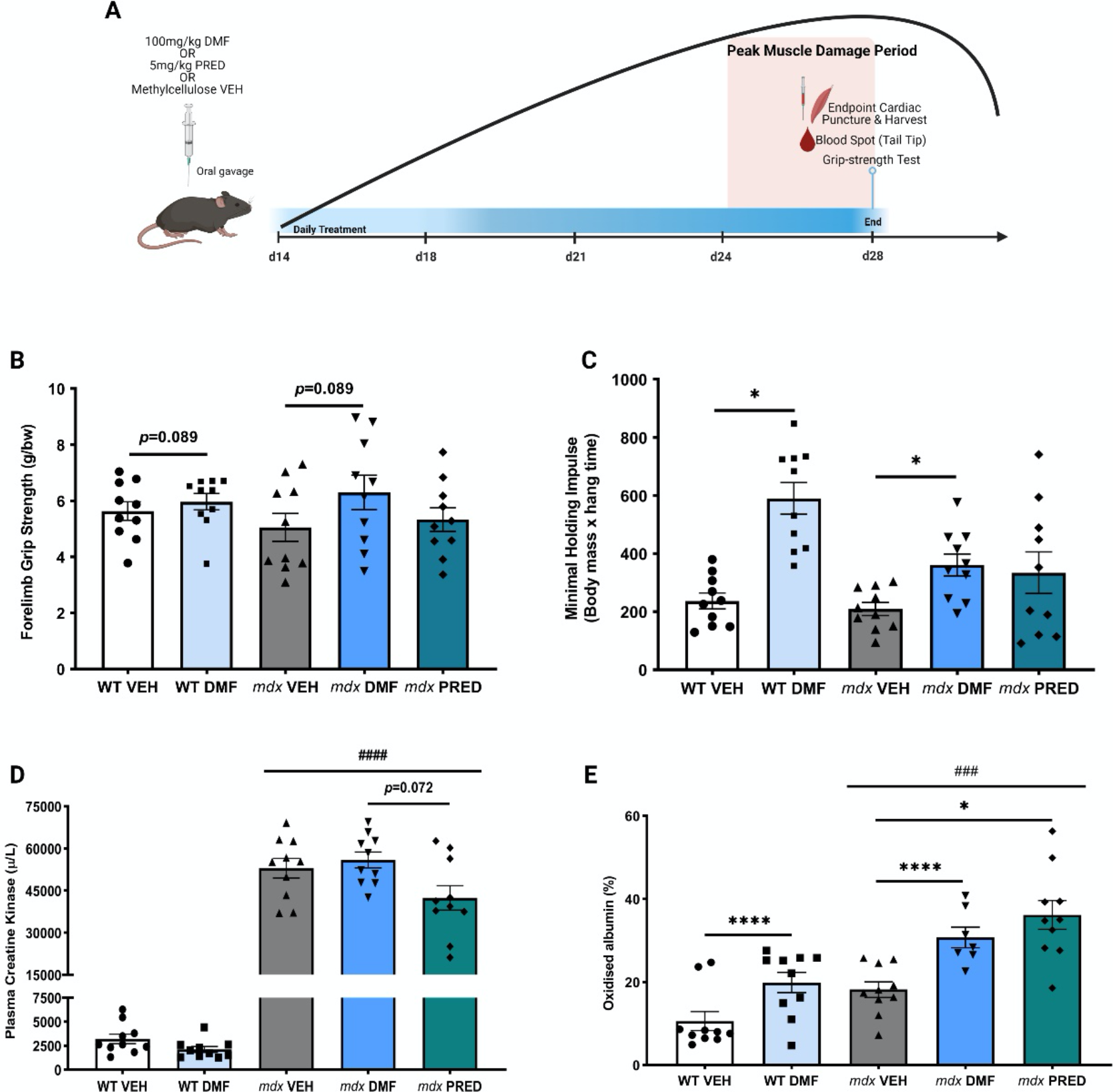
Dimethyl fumarate (DMF) improves muscle function but not Duchenne Muscular Dystrophy (DMD) blood biomarkers. (A) Schematic of the treatment period and clinically compatible testing protocol beginning at 14 days of age and ceasing at 28 days of age. Mice were treated daily via oral gavage with either vehicle (0.5% methylcellulose; VEH), 100mg/kg DMF or 5mg/kg prednisone (PRED) and underwent grip strength and blood biomarker testing at the experimental endpoint at 28 days of age. B) Forelimb and (C) whole-body grip strength was comparable between wild-type (WT) and *mdx* VEH mice at 28 days yet DMF increased grip strength in *mdx* (and WT) mice (trend for forelimb strength, significant for whole body holding impulse). (D) Creatine kinase (CK) levels were significantly elevated in *mdx* mice compared to WT mice but were not improved by either DMF or PRED treatment, although PRED treatment trended to reduce CK levels compared to DMF treated *mdx* mice. (E) Blood oxidised albumin levels were comparable between *mdx* and WT mice at 28 days of age. Both DMF and PRED stimulated albumin oxidisation in both WT (DMF only) and *mdx* (DMF and PRED) mice. Treatment effect: \**p*<0.05, \*\*\*\**p*<0.0001; genotype effect: ^###^*p*<0.001

Despite juvenile growth inducing acute severe muscle damage in *mdx* mice sufficient to raise the haematologic clinical biomarker, creatine kinase (CK), TREAT-NMD reference data report stable functional strength testing until ~5 weeks of age after which strength decline becomes evident (21). Consistent with the reference, *mdx* mice showed stable forelimb (Fig 2B) and whole body (Fig 2C) grip strength yet ~24-fold elevated plasma CK levels at endpoint (compared to wild-type (WT); Fig 2D). Neither DMF nor PRED lowered plasma CK levels at endpoint compared to *mdx* VEH (Fig 2D) yet there was a trend for DMF to increase maximal weight-corrected grip strength (treatment effect, *p*=0.085) and it significantly increased the maximum holding impulse derived from the whole-body hang test in WT and *mdx* mice (Fig 2C). PRED had no effect on muscle function tests.

Novel blood biofluid biomarkers of DMD progression are currently being validated in patients (22, 23) and animal models (24, 25). In this study, we assessed albumin oxidation as a biomarker of systemic oxidative stress. Albumin is the major haematologic antioxidant and exerts both functional and protective roles in the presence of oxidative stress (26). Increased albumin oxidation was observed in *mdx* compared to WT blood at endpoint (Fig 2D, genotype effect), the theoretical “peak” of the juvenile muscle damage period. Unexpectedly, both DMF and PRED increased blood albumin oxidation (including in WT mice for DMF). Previous studies in patients with RRMS demonstrated DMF to transiently elevate the oxidative state of peripheral blood by driving reactive oxygen species (ROS) production in monocytes (27). This “oxidative burst” appears crucial for DMF’s immunomodulatory precision. To our knowledge, these data are the first to demonstrate that PRED acts in a similar fashion.

### DMF activates Nrf2 and the cytoprotective and anti-inflammatory program in skeletal muscle

Constitutive Nrf2 synthesis outside of Keap1’s control can result in higher Nrf2 expression, although this is not necessary for effective signalling of the Phase II antioxidant response. We assessed protein levels and activation of key regulators of Nrf2 via western immunoblot. Our data show DMF successfully targets Nrf2 in skeletal muscle (gastrocnemius) with a tendency to increase Nrf2 levels (*p*=0.07 treatment effect; Fig 3A) leading to the significant upregulation of key Phase II antioxidant enzymes including NAD(P)H dehydrogenase: quinone oxidoreductase (NQO1; WT and *mdx* DMF versus VEH; Fig 3B) and superoxide dismutase 1 (SOD1; Fig 3C). Haemoxygenase 1 (HO-1), a strong suppressor of ROS and inflammation, was already elevated in *mdx* muscle and DMF further induced upregulation in *mdx* but not WT muscles (*p*=0.059 *mdx* DMF versus *mdx* VEH; Fig 3D). The expression of Keap1 (Fig 3E), as well as phosphorylated p62 (sequestosome 1; Supp Fig 2A), a classical receptor of autophagy which sequesters and tags Keap1 for degradation, were also elevated in *mdx* compared to WT muscle. DMF treatment increased p62 protein expression in WT and *mdx* muscle (Fig 3F) and tended to increase phosphorylation of the p62^Ser349^ residue, which maintains Keap1 binding and Nrf2 activity (*p*=0.058 treatment effect; Supp Fig 2A). Neither the activity (oxidised Cys^106^), nor the protein expression of Nrf2’s molecular chaperone, DJ-1 (also known as Parkinson’s disease protein 7 (PARK7)), were affected by DMF treatment although DJ-1 protein expression was reduced in *mdx* compared to WT muscle (Supp Fig 2B-C). These data suggest (1) oxidative stress is already present in juvenile *mdx* muscles sufficient to induce adaptive hormesis; and (2) DMF can augment the myocellular control of hormesis through activating Nrf2.

**Figure 3:**
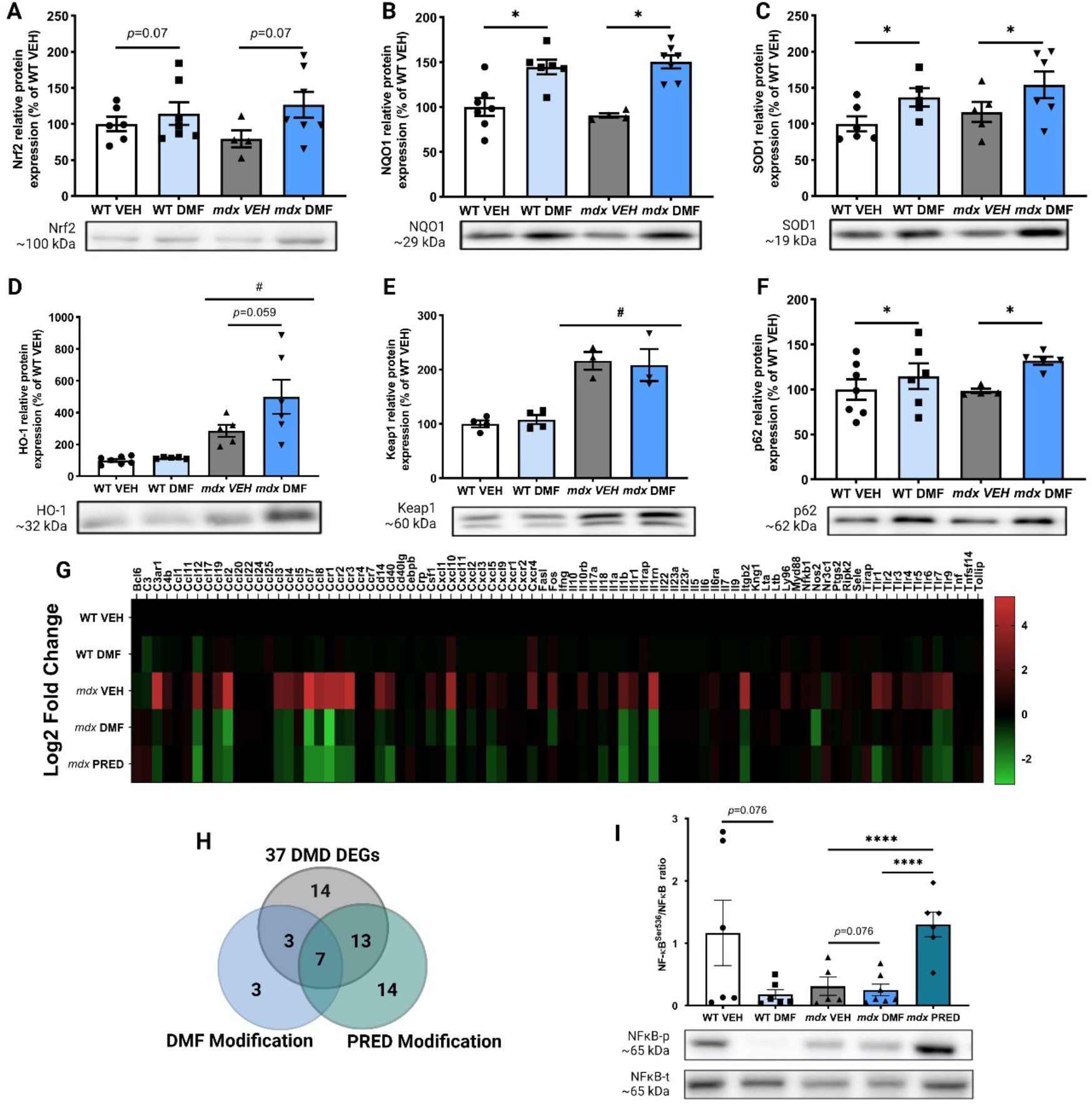
Dimethyl fumarate (DMF) activates nuclear erythroid 2-related factor 2 (Nrf2) and induces the Phase II antioxidant response in *mdx* skeletal muscle. (A) There was a strong trend for Nrf2 protein to increase due to DMF treatment in wild-type (WT) and *mdx* muscle. DMF treatment increased the expression of the key downstream Phase II antioxidants, NAD(P)H dehydrogenase: quinone oxidoreductase (NQO1; B) and superoxide dismutase 1 (SOD1; C). (D) Hemeoxygenase-1 (HO-1) was elevated in *mdx* mice compared to WT mice, in which there was a strong trend for DMF treatment to further increase HO-1 expression. (E) Keap1 protein was elevated in *mdx* but DMF had no effect. (F) Total p62 protein expression was comparable between WT and *mdx* muscle with DMF inducing further p62 expression in both strains. (G-H) An array of inflammatory genes was assessed. In the *mdx* quadriceps, 37 differentially expressed genes (DEGs) compared to WT were detected, in which DMF treatment modified 3 DEGs, while prednisone (PRED) modified 13 DEGs. (I) There was a strong trend for ratio of phosphorylated nuclear factor kappa B (NF-κB) to total NF-κB to be reduced following DMF treatment in both WT and *mdx* muscles whilst PRED had the opposite effect, elevating the ratio significantly compared to both vehicle (VEH) and DMF treated *mdx* mice. Treatment effect: \**p*<0.05, \*\*\*\**p*<0.0001; genotype effect: ^#^*p*<0.05.

As well as anti-oxidation, DMF also functions as a potent anti-inflammatory drug (Fig 1). PRED is also a potent immunosuppressant to confer disease modifying benefit in DMD. To test the immunomodulatory capacity of DMF compared to PRED, we profiled 84 inflammatory genes via qPCR RT2 gene array in gastrocnemius muscles. 37 (45%) genes were differentially regulated in juvenile *mdx* compared to WT muscle (Fig 3G-H; Table 1). Most were associated with the acute-phase response or general regulation of inflammation. Ten differentially expressed genes (DEGs) were increased by >10-fold, which were typically chemokines or chemokine/cytokine receptors (Table 1). Consistent with its known immunosuppressant action, PRED downregulated a more extensive inflammatory gene profile than DMF (54% versus 27% of *mdx* DEGs) and this was generally by the magnitude of 2-4-fold. Although modulating fewer genes, DMF downregulated the expression of key inflammatory genes by a much larger magnitude than PRED, e.g., DMF downregulated the gene expression of chemokine, CCL7, and CCR1, the type-1 receptor for chemokines, CCL3, CCL5, CCL7 and CCL23, by 7- and 10-fold, respectively (versus non-significantly and 6-fold for PRED, respectively). CCL7 was the most DEG in *mdx* muscle (increased by 35-fold compared to WT VEH) and is of particular significance since it interacts with matrix metalloproteinase 2 (MMP2), a recently identified seed gene that drives the DMD disease module (28) via chemokine receptor 2 (CCR2) (29). Dysregulated CCL7 is implicated in, and worsens, immunological diseases including MS (30) and psoriasis (31) against which DMF is particularly effective. DMF also had fewer off-target modulatory effects on normally regulated genes (NEGs) than PRED (3 versus 6 inflammatory NEGs, respectively). Our data importantly show that DMF more selectively modulates the inflammation program that drives the DMD disease module in juvenile *mdx* mice.

**Table 1:** Effect of dimethyl fumarate (DMF) versus prednisone (PRED) treatment on differentially expressed genes (DEGs) involved in inflammation in *mdx* compared to wild-type (WT) muscle (in order of most to least dysregulated). DEGs criteria are >2-fold regulation, *p*<0.05 from reference group (*mdx* VEH v WT VEH). Bolded font indicates significantly modulated by drug (any fold change; p<0.05). Key: AP: acute phase; CM: cytokine metabolism: CMS: cytokine-mediated signalling; CR: chronic response; IRR: inflammatory response regulation; \**p*<0.05, \*\**p*<0.01, \*\*\**p*<0.001, \*\*\*\**p*<0.0001.

| DEG (*mdx* v *WT*) |  | DMF Benefit |  | PRED Benefit |  | Response Type | Other Functions |
| --- | --- | --- | --- | --- | --- | --- | --- |
| Gene | Fold Reg. | Fold Reg. | $p$ | Fold Reg. | $p$ | | |
| *CCL7* | ↑ 34.57 | ↓ **7.06** | * | ↓ 4.84 | 0.167 | IRR | +CR |
| *CCL2* | ↑ 25.56 | ↓ 5.55 | 0.071 | ↓ 3.21 | 0.274 |  | Humoral |
| *CCR3* | ↑ 24.00 | ↓ **1.86** | * | ↓ **3.04** | ** |  | CMS |
| *C3AR1* | ↑ 22.40 | ↓ 1.59 | 0.101 | ↓ 2.24 | * |  |  |
| *CCR1* | ↑ 17.48 | ↓ **10.18** | *** | ↓ **6.03** | * | AP | CMS |
| *CCR2* | ↑ 16.23 | ↓ 1.65 | 0.097 | ↓ **3.10** | * | Humoral, CMS |  |
| *IL1RN* | ↑ 15.83 | ↓ 4.72 | 0.406 | ↓ **4.37** | * | AP | CMS |
| *CCL8* | ↑ 13.15 | ↓ **1.82** | * | ↓ **4.65** | ** | IRR |  |
| *ITGβ2* | ↑ 12.86 | ↓ 2.01 | 0.103 | ↓ 2.66 | 0.057 |  | Humoral |
| *CXCL10* | ↑ 12.13 | ↓ **3.13** | * | ↓ **2.27** | ** |  |  |
| *CCL12* | ↑ 6.95 | ↓ 3.03 | 0.141 | ↓ **4.74** | * |  |  |
| *CD14* | ↑ 6.69 | ↓ 1.75 | 0.103 | ↓ **1.99** | * |  |  |
| *CCL3* | ↑ 5.41 | ↓ 2.06 | 0.100 | ↓ 2.05 | 0.412 |  | Humoral |
| *TLR9* | ↑ 5.18 | ↓ **2.06** | ** | ↓ **2.78** | ** |  |  |
| *TLR1* | ↑ 5.15 | ↓ 1.31 | 0.344 | ↓ **3.09** | * |  | CM |
| *CCL4* | ↑ 4.96 | ↓ 2.29 | 0.100 | ↓ 2.26 | 0.160 | IRR |  |
| *IL1β* | ↑ 4.44 | ↓ 3.4 | 0.101 | ↓ 4.11 | 0.062 | AP | +CR,<br>humoral |
| *CXCR4* | ↑ 4.32 | ↓ 1.17 | 0.274 | ↓ 1.58 | 0.053 | IRR |  |
| *TLR7* | ↑ 3.66 | ↓ 2.33 | 0.094 | ↓ **2.39** | * |  |  |
| *CD40* | ↑ 3.24 | ↓ 1.39 | 0.01 | ↓ **3.36** | ** |  |  |
| *TLR2* | ↑ 3.40 | ↓ 1.18 | 0.121 | ↓ **1.57** | ** |  |  |
| *CCL19* | ↑ 3.15 | ↓ **2.56** | ** | ↓ **2.57** | * |  |  |
| *CCL5* | ↑ 3.10 | ↑ 1.27 | 0.215 | ↓ **1.48** | ** | AP | +CR,<br>chemokine |
| *FOS* | ↑ 3.51 | ↓ **2.22** | * | ↑ 1.32 | 0.393 | IRR |  |
| *IL1R1* | ↑ 2.56 | ↓ **2.02** | * | ↓ **2.04** | ** |  | CMS |
| *IL10Rβ* | ↑ 2.55 | ↓ 1.3 | 0.113 | ↓ 1.26 | 0.167 |  |  |
| *TLR6* | ↑ 2.47 | ↓ 1.15 | 0.462 | ↓ **1.76** | * |  | CM |
| *LY96* | ↑ 2.40 | ↓ 1.08 | 0.231 | ↓ 1.2 | 0.140 |  | Humoral |
| *TLR4* | ↑ 2.26 | ↓ 1.1 | 0.544 | ↓ 1.19 | 0.139 |  | CM |
| *CSF1* | ↑ 2.18 | ↓ **1.77** | * | ↓ 1.22 | 0.329 |  |  |
| *C4B* | ↑ 2.13 | ↑ 1.17 | 0.330 | ↓ 1.01 | 0.673 |  | Humoral |
| *TLR5* | ↑ 1.88 | ↓ 1.15 | 0.462 | ↓ **1.76** | * | IRR |  |
| *IL18* | ↑ 1.77 | ↓ 1.31 | 0.218 | ↓ 1.88 | 0.073 |  | CM |
| *NR3C1* | ↓ 1.76 | ↑ 1.19 | 0.296 | ↑ **1.86** | * | IRR |  |
| *IL6Ra* | ↑ 1.58 | ↑ 1.11 | 0.665 | ↑ 1.35 | 0.225 |  | CMS |
| *C3* | ↓ 1.57 | ↑ 1.23 | 0.257 | ↑ 1.66 | 0.185 | IRR | Humoral |
| *BCL6* | ↓ 1.50 | ↑ 1.29 | 0.149 | ↑ **1.76** | * | IRR |  |

We also assessed activation and protein expression of the master regulator of innate immunity, nuclear factor kappa B (NF-κB), which controls a complex signaling hub linking pathogenic signals with cellular danger signals. NF-κB is purportedly suppressed by both Nrf2 and PRED and has also been linked to control of mitochondrial respiration (32). Total NF-κB protein levels were equivalent between *mdx* and WT mice and neither DMF nor PRED treatment modulated them (Supp Fig 2D). Phosphorylation of the Ser^536^ activation site was also equivalent between WT and *mdx* VEH muscles (there was high variability especially in WT control muscles; Supp Fig 2E), suggesting NF-κB signalling is crucial for muscle growth and re-modelling in juvenile mice, though no more active due to dystrophin-deficiency. Most surprisingly, PRED treatment increased phosphorylated (Ser^536^) NF-κB expression higher than in any other group (*mdx* PRED versus all other groups) whereas DMF tended to lower levels in both WT and *mdx* muscles (*p*=0.058 treatment effect, Supp Fig 2E). These data were mimicked in the ratio of Ser^536^ NF-κB phosphorylation to the total NF-κB protein, a biomarker of NF-κB activity (Fig 3I) and demonstrate DMF and PRED confer different anti-inflammatory actions on *mdx* skeletal muscle. The significance of these data for the progression of DMD is unknown since: (1) the anti-inflammatory and immunomodulatory effects of PRED are well known and disease modifying; and (2) NF-κB signaling is complex and may elicit both beneficial and negative effects depending upon the biological context.

### DMF improves force production and protects against fatigue in fast-twitch muscle

Muscle force production relative to mass is predictive of muscle quality and perhaps the most useful indicator of drug benefit on DMD progression (33). Since there was no impact of juvenile disease on whole-body functional force measures in *mdx* mice (Fig 2B), we studied *ex vivo* contractile characteristics at the muscle level for early signs of function anomalies. We assessed predominantly type II extensor digitorum longus (EDL) and type I soleus (SOL) muscles to scope for fibre-type specific effects of DMF and PRED (data presented in Table 2). The specific (cross sectional area (CSA)-corrected; sPo) force was ~70% lower in EDL (*mdx* compared to WT VEH; Fig 4A) and ~30% lower in SOL (*mdx* compared to WT VEH; Fig 4B). DMF and PRED improved the specific force of EDL by >3-fold (Fig 4A) but had no effect on SOL (Fig 4B), suggesting specific effects on fast type II muscle fibres. However, DMF shifted the force-frequency curve of WT and *mdx* VEH EDL and SOL muscles to the left (treatment effect, Fig 4C-D) indicating pan modulation of cross-bridge sensitivity irrespective of fibre type. EDL and SOL muscles were subsequently subjected to a fatigue protocol and muscle force production shown at minute intervals (Fig 4E-F; *p*<0.001 fatigue effect for EDL and SOL). DMF, but not PRED, protected *mdx* EDL muscles from fatigue (treatment effect; Fig 4E). *Mdx* SOL muscles fatigued much less during the protocol (since they produced such little force in response to stimuli and due to their predominantly type I oxidative phenotype, e.g., see Table 2). Neither DMF nor PRED affected fatigability of SOL (Fig 4F).

**Table 2:** Contractile function characteristics of extensor digitorum longus (EDL) and soleus muscle from wild-type (WT) and *mdx* mice treated with vehicle (VEH), dimethyl fumarate (DMF) or prednisone (PRED). Significance: ^##^*p*<0.01, ^###^*p*<0.001, ^####^*p*<0.0001 (*mdx* VEH v WT VEH); \**p*<0.05, \*\**p*<0.01, \*\*\*\**p*<0.0001 (VEH v DMF); ^*p*<0.05, ^^^*p*<0.001 (DMF v PRED).

|  | EDL |  |  |  |  | Soleus |  |  |  |  |
| --- | --- | --- | --- | --- | --- | --- | --- | --- | --- | --- |
|  | WT |  | *mdx* |  |  | WT |  | *mdx* |  |  |
|  | VEH | DMF | VEH | DMF | PRED | VEH | DMF | VEH | DMF | PRED |
| Muscle mass (mg) | 4.98 ±<br>0.51 | 4.32 ±<br>0.37 | 4.39 ±<br>0.11 | 4.45 ±<br>0.28 | 5.78 ±<br>0.47 | 5.95 ±<br>0.56 | 4.78 ±<br>0.36** | 6.46 ±<br>0.41 | 3.59 ±<br>0.55** | 6.34 ±<br>0.15^ |
| Optimum length (L<sub>o</sub>; mm) | 7.9 ±<br>0.57 | 5.57 ±<br>0.19**** | 9.8 ±<br>0.20## | 5.75 ±<br>0.34**** | 9.41 ±<br>1.02^^^ | 7.00 ±<br>0.63 | 4.88 ±<br>0.21**** | 10.60 ±<br>0.10### | 4.79 ±<br>0.20**** | 8.42 ±<br>0.80^^^ |
| Absolute (maximal isometric)<br>force (mN) | 147.11<br>± 27.42 | 81.02 ±<br>14.11 | 30.73 ±<br>6.05## | 83.45 ±<br>13.10## | 92.05 ±<br>19.10 | 100.31 ±<br>21.80 | 74.09 ±<br>15.53 | 35.88 ±<br>5.66# | 19.27 ±<br>5.65# | 45.58 ±<br>8.64 |

**Figure 4:**
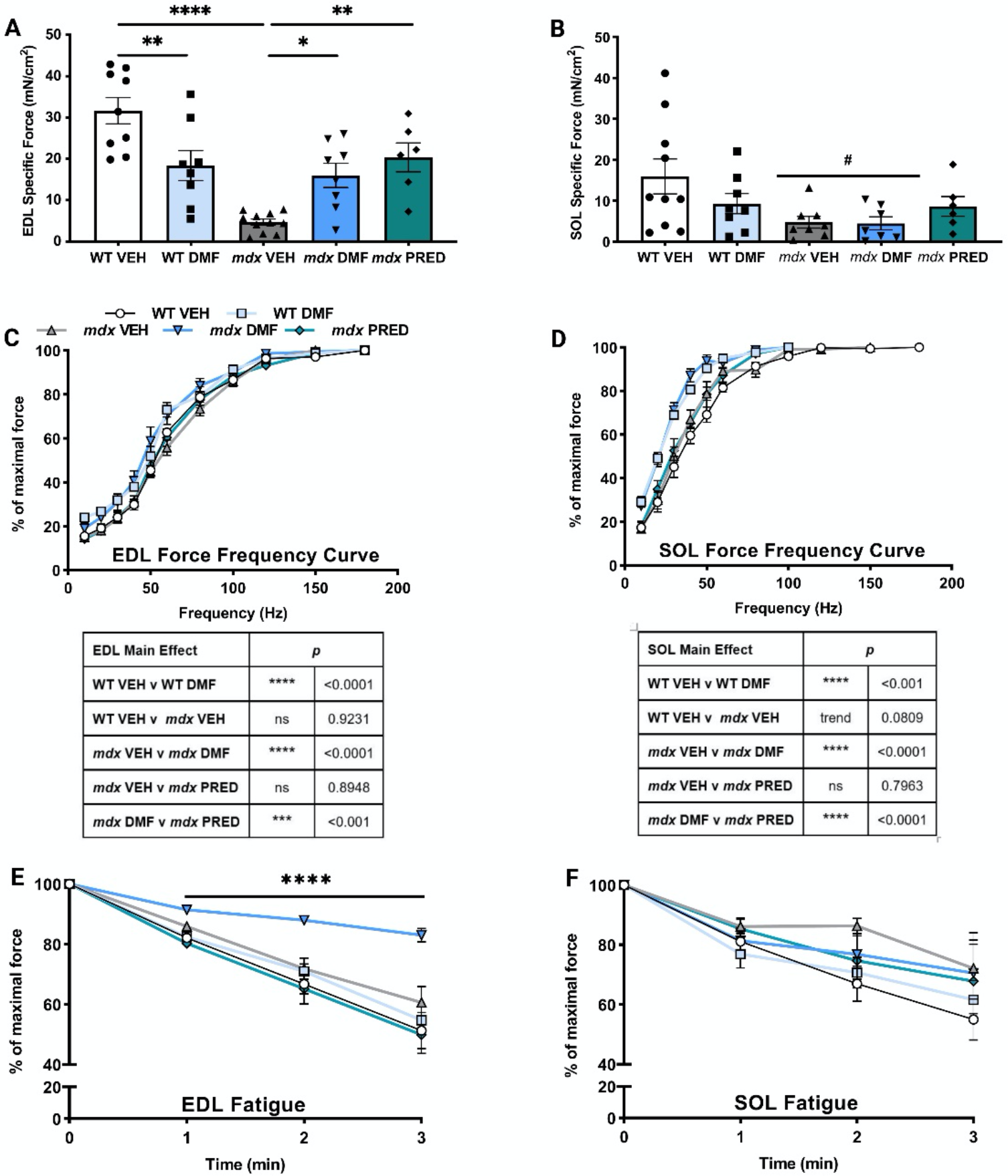
Dimethyl fumarate (DMF) recovers force and reduces the fatiguability of type II *mdx* extensor digitorum longus (EDL) muscles. (A) The specific force of EDL was significantly lower in *mdx* vehicle (VEH) compared to wild-type (WT) and DMF treatment induced opposite effects in WT and *mdx* mice. In WT EDL, DMF reduced the specific force, while DMF significantly increased EDL specific force in *mdx* mice. Prednisone (PRED) also increased the specific force of *mdx* EDL. (B) As with EDL, soleus (SOL) specific force was lower in *mdx* mice. However, neither DMF nor PRED treatment were ameliorative. DMF treatment shifted the force-frequency curve to the left in both WT and *mdx* EDL (C) and SOL (D) muscles. (E) DMF also improved the fatiguability of the *mdx* EDL but had no effect in the SOL (F). Treatment effect: \**p*<0.05, \*\**p*<0.01, \*\*\*\**p*<0.0001; genotype effect: ^#^*p*<0.05

### DMF augments mitochondrial respiration in *mdx* fibres through anaplerosis

Channelling fumarate through the malate-aspartate shuttle into the mitochondrial Kreb’s cycle can reverse flux, driving mitochondrial respiration through Complex II/ succinate dehydrogenase (SDH). This mechanism is used endogenously when the purine nucleotide cycle is activated by degrading purines during metabolic stress, which generates fumarate for mitochondrial oxidative phosphorylation augmentation (34). Mitochondrial biogenesis and networking (i.e., fission and fusion dynamics), as well as the biosynthesis of purines, are also controlled by Nrf2 activity (14). Mitochondrial respiratory function was measured in flexor digitorum brevis (FDB) fibres using extracellular flux and a mitochondrial stress test involving the sequential application of inhibitor/stimulator drugs (Fig 5A). There was a trend for mitochondrial oxygen consumption in the basal and phosphorylating states to decrease in *mdx* compared to WT VEH FDB fibres (data not shown), consistent with previous studies in older mice (35). However, this effect was abolished by correction for protein concentration (Fig 5A-B). Only non-mitochondrial respiration, which is mostly attributed to cellular oxidase activity associated with anti-inflammation and antioxidation, was reduced in mdx FDB fibres (Fig 5A-B). Nevertheless, DMF increased the basal, phosphorylating, maximal and non-mitochondrial protein-corrected respiration (Fig 5A-B) in *mdx* FDB fibres resulting in an overall higher bioenergetical state (Fig 5C). There was no evidence of mitochondrial uncoupling in response to increased DMF-dependent substrate flux (Fig 5D). Spare reserve capacity (SRC) is an internally normalised determinant of mitochondrial fitness/flexibility and depends on electron transport chain (ETC) and inner membrane integrity, bioenergetical demand and preservation of mitochondrial homeostasis. These factors are controlled by several signalling pathways associated with Nrf2 including cytokine mediated STAT3 signalling, glucose and fatty acid metabolism and oxidative stress (see (37) for excellent review). DMF increased the SRC in WT and *mdx* muscles consistent with anaplerosis of the TCA cycle (Fig 5E).

**Figure 5:**
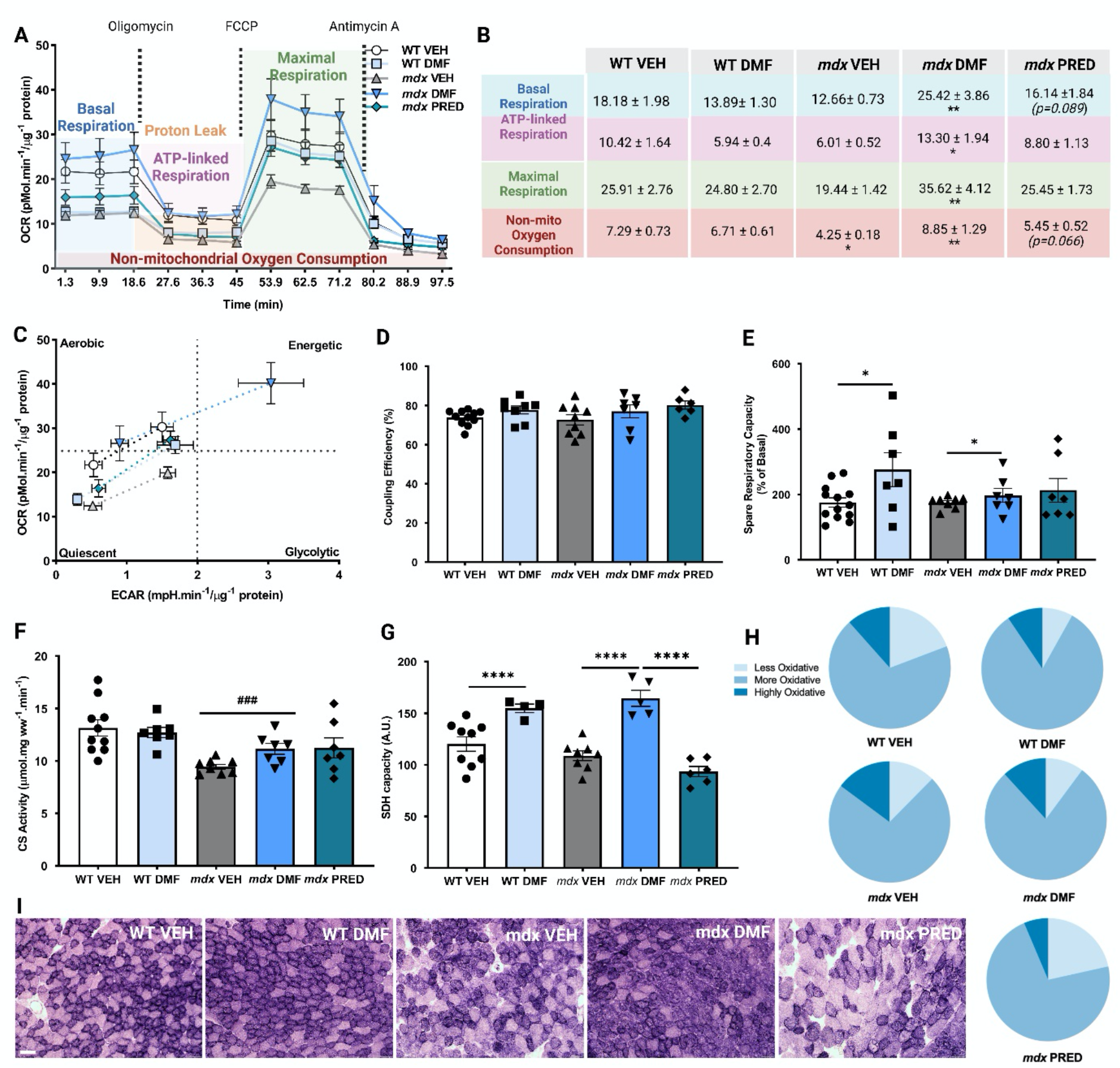
Dimethyl fumarate (DMF) enhances mitochondrial respiratory function, including basal, ATP production-associated and maximal respiration in *mdx* flexor digitorum brevis (FDB) fibres. (A-B) In *mdx* fibres, non-mitochondrial oxygen respiration was lower compared to wild-type (WT). DMF treatment increased basal, ATP-linked, maximal, and non-mitochondrial oxygen consumption in *mdx* fibres only. There was a strong trend for prednisone (PRED) treatment to increase basal respiration and non-mitochondrial oxygen consumption in *mdx* mice, but not other respiration parameters. The overall gain in bioenergetical function induced by DMF is observed in (C). (D) Coupling efficiency was comparable across groups, (E) Spare respiratory capacity was not affected by genotype, but DMF increased it in both WT and *mdx* FDB fibres. (F) Citrate synthase (CS) activity was reduced in *mdx* muscles but was not modulated by DMF or PRED treatment. (G) DMF did, however, increase succinate dehydrogenase (SDH) capacity in both WT and *mdx* muscle, whereas PRED had no effect. (H) The proportion of less, more, and highly oxidative muscle fibres is shown. (I) Representative images of the SDH stain. Treatment effect: \**p*<0.05, \*\**p*<0.01, \*\*\*\**p*<0.0001; genotype effect: ^###^*p*<0.001.

Citrate synthase (CS) activity, a classical biomarker of mitochondrial content, was reduced in *mdx* VEH gastrocnemius but was not modulated by DMF or PRED (Fig 5F), nor were protein biomarkers of biogenesis (TFAM, PGC1α), fission (DRP-1) or fusion (OPA-1) signalling (Supp Figure 3A-D). Since DMF (and PRED’s) protective effects on muscle force production were fibre type specific, we also assessed Complex II/SDH capacity in predominantly fast-twitch tibialis anterior (TA) sections. Consistent with Kreb’s reversal and increased flux of malate>fumarate>succinate through Complex II, DMF increased SDH capacity in both WT and *mdx* muscle (Fig 5G). Based on SDH activity staining, DMF drove a more oxidative phenotype while PRED drove a less oxidative phenotype (compared to VEH; Fig 5H-I and Supp Figure 3F-H) demonstrating stark differences between these drugs on metabolic plasticity. Our data highlight pro-mitochondrial effects of DMF induced by fumarate-stimulated anaplerosis rather than via improved mitostasis (e.g., biogenesis, fission-fusion dynamics, efficiency), which accounts for the fatigue resistance conveyed on fast-twitch muscle (Fig 4E). DMF could be useful to treat the known mitochondrial dysfunction in muscle from older *mdx* mice and DMD patients as the disease progresses.

### DMF modifies biomarkers of muscle integrity, quality, and histopathology

To test whether DMF-induced Nrf2 activation could improve histopathology, a subset of juvenile WT and *mdx* mice were injected with Evan’s blue dye (EBD), a cell-impermeant extravasation dye that can only be absorbed by muscle fibres with membrane damage. EDL, SOL, TA, and diaphragm (DIA) muscles were collected 24h post-EBD injection. DMF treatment significantly reduced the % of EBD positive fibres in all *mdx* muscles by up to 6-fold (Fig 6A-D). In contrast, PRED had no effect on hindlimb muscles (Fig 6A-C) but was just as effective as DMF at reducing membrane damage of DIA muscles (Fig 6D). Nrf2 activator compounds, sulforaphane and curcumin, also reduce EBD fluorescence in *mdx* muscle fibres (16, 19) while DMF specifically tempers dysregulated phospho- and sphingo-lipid metabolism by inhibiting damaging lipases (38), highlighting membrane protection is mediated via Nrf2.

**Figure 6:**
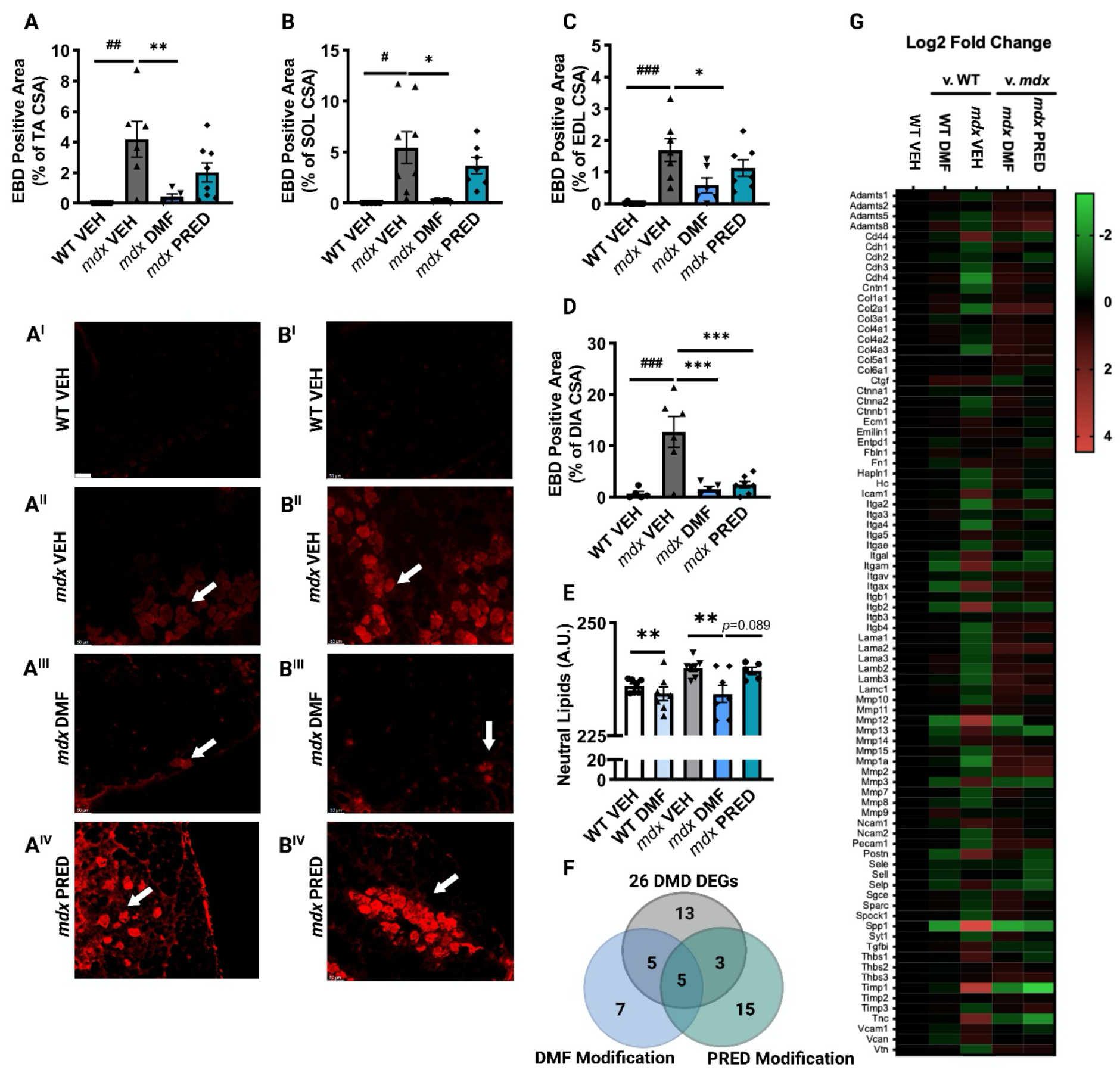
Dimethyl fumarate (DMF) improves biomarkers of muscle pathology. (A-D) Evans blue dye (EBD) permeation into muscle fibres, a biomarker of compromised sarcolemma integrity, was increased in the *mdx* tibialis anterior (TA), soleus (SOL), extensor digitorum longus (EDL) and diaphragm (DIA). DMF reduced the EBD positive area of all muscles. Comparatively, prednisone (PRED) reduced EBD fluorescence in DIA only. (E) Muscle neutral lipids, as quantified by Oil Red O staining, were decreased following DMF treatment in WT and *mdx* TA. (F-G) An array of extracellular matrix genes was assessed. In *mdx* compared to WT quadriceps, 26 differentially expressed genes (DEGs) were detected. DMF treatment modified 10 DEGs, while PRED modified 8 DEGs – 5 DEGS were modified by both drugs. Treatment effect: \**p*<0.05, \*\**p*<0.01, \*\*\**p*<0.001; genotype effect: ^#^*p*<0.05, ^##^*p*<0.01, ^###^*p*<0.001.

In DMD patients, muscle quality is severely compromised. As well as having less muscle due to chronic fibre degeneration, fibro/adipogenic progenitors within the ECM drive reactive steatosis (fat infiltration) and/or fibrosis in response to persistent sterile inflammation signals – all three factors contribute to progressive loss of function (39) (for review of biology see (40)). DMD muscles from patients and *mdx* mice also retain more intracellular lipids, presumably due to the extensive mitochondrial and metabolic perturbations characteristic of the disease, which leach from myofibres and contribute to fatty replacement of muscle (34). Although our mice were too young to observe extensive steatosis and fibrosis (which is relatively mild in *mdx* mice compared to DMD patients anyway), we assessed early signs via (1) Oil Red O (ORO) staining of neutral lipids in TA sections from non-EBD-injected mice; and (2) qPCR RT2 gene array of genes controlling ECM composition and cell adhesion. DMF reduced myocellular lipid content in both WT and *mdx* muscles (Fig 6E) consistent with augmentation of lipid metabolism (38) and mitochondrial function (Fig 5). 26 ECM genes were differentially expressed by >1.5-fold (*p*<0.05) in *mdx* compared to WT (VEH) muscle (see Fig 6F-G and Table 3) demonstrating activation of a complex re-modelling program in juvenile mice undergoing an acute disease phase. Of these DEGs, DMF modulated 10 (38%) while PRED modulated 8 (31%) – 5 of the same genes were modulated by both drugs. 2 DEGs, *SPP1* (secreted phosphoprotein 1) and *TIMP1* (tissue inhibitor of metalloproteinase 1), were upregulated by >10-fold, reversibly modulated by both DMF and PRED (*SPP1* more so by DMF and *TIMP1* more so by PRED; Table 3) and in addition to MMP2, were also defined as seed genes within the DMD disease module (28). *SPP1* encodes osteopontin (OPN) whose ablation lessens severity of the *mdx* phenotype by skewing macrophage polarisation to drive a pro-regenerative over a pro-fibrogenic phenotype (41). Importantly, gene expression of MMP12 (macrophage elastase) was >6-fold higher in *mdx* compared to WT VEH muscle and DMF, but not PRED, reduced expression by >2-fold. MMP12 is induced by macrophage activity and degrades the ECM resulting in loss of muscle elasticity (42). Interestingly, DMF also normalised the longer optimum length (L_o_) observed in *mdx* compared to WT (VEH) EDL and SOL muscles during our contractile studies (Table 2) although the long-term significance of this is unknown since DMF improved contractile function in *mdx* EDL but not SOL muscle. Although fibrosis-associated gene, *TGFBI* (transforming growth factor β inducible), just fell short of the 1.5-fold DEG classification cut off, its expression was significantly increased in *mdx* VEH compared to WT VEH muscle (1.42-fold increase, *p*<0.01) suggesting induction of the fibrosis program even in juvenile mice. DMF and PRED effectively reduced *TGFBI* expression (both by 1.6-fold, *p*<0.001) highlighting comparable efficacy in suppressing the fibrosis gene program.

**Table 3:**
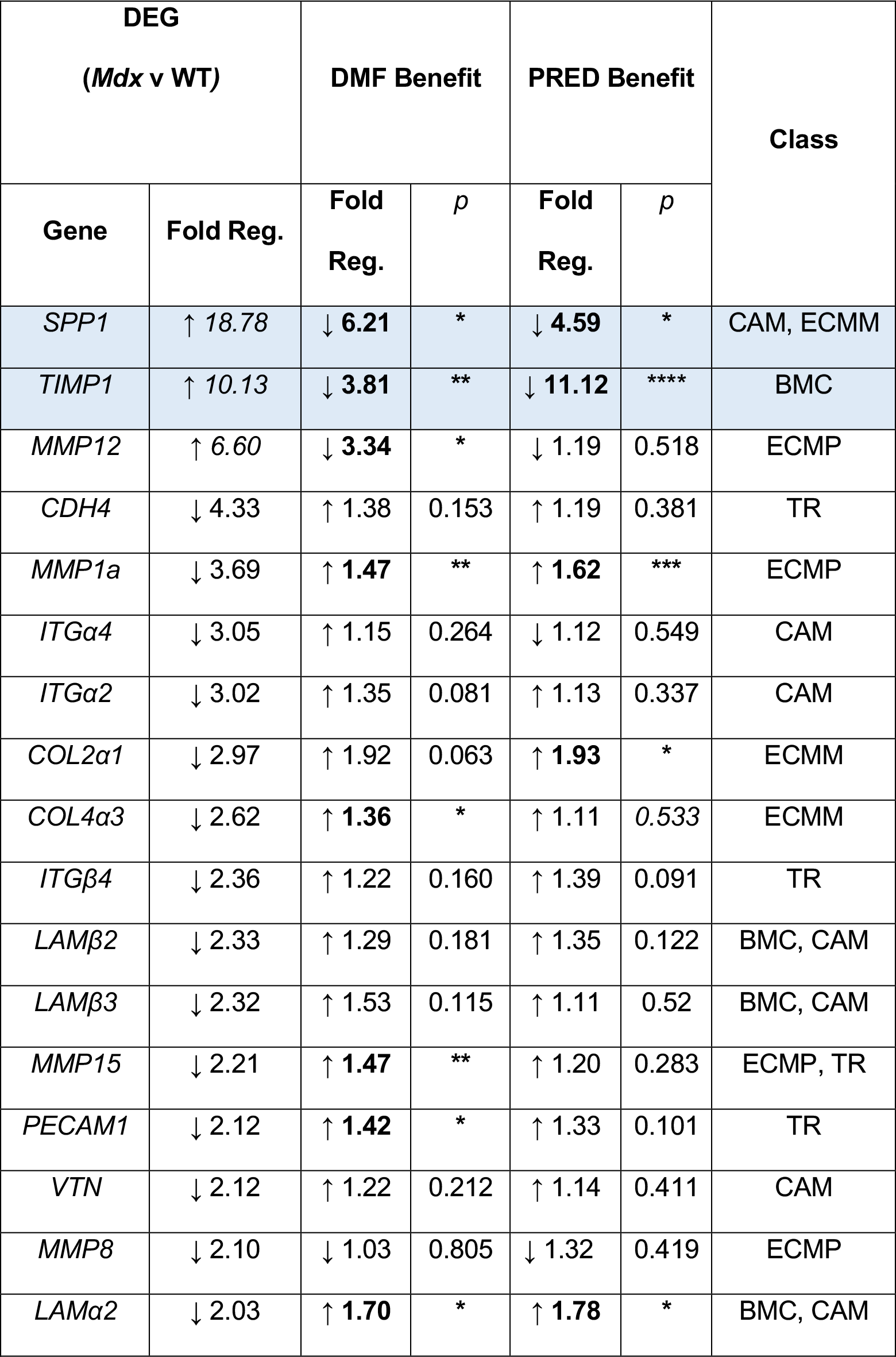
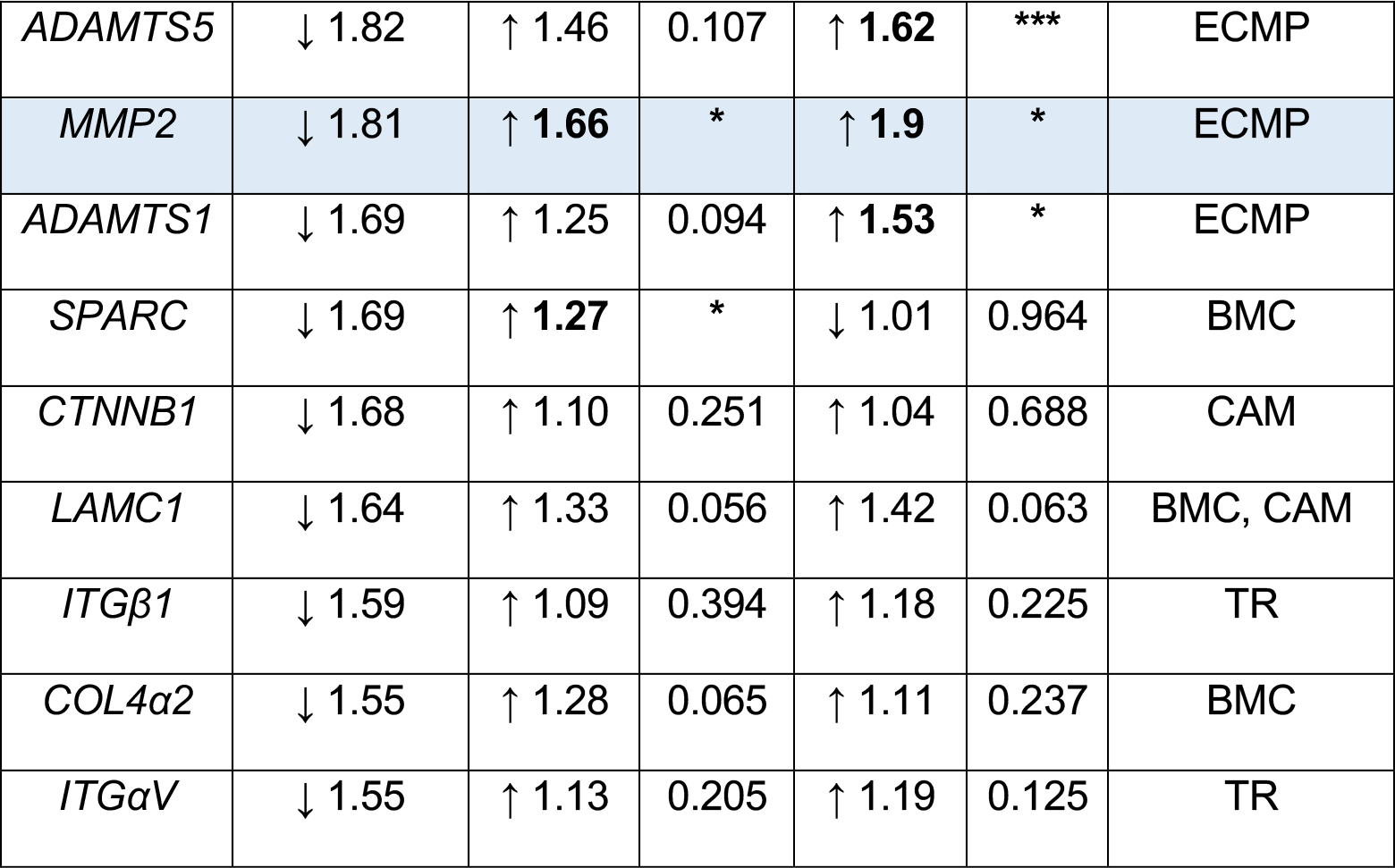
Effect of dimethyl fumarate (DMF) versus prednisone (PRED) treatment on differentially expressed genes (DEGs) involved in extracellular matrix composition and cell adhesion in *mdx* compared to wild-type (WT) muscle (in order of most to least dysregulated). DEGs criteria are >2-fold regulation, *p*<0.05 from reference groups (WT VEH v *mdx* VEH). Bolded font indicates significantly modulated by drug (any fold change; *p*<0.05). Key: BMC: basement membrane constituents; CAM: cell adhesion molecule; ECMM: extracellular matrix molecules; ECMP: extracellular matrix proteases; TR: transmembrane receptors: \**p*<0.05, \*\**p*<0.01, \*\**p*<0.001, \*\*\*\**p*<0.0001. Blue shading indicates DMD disease seed genes.

As well as modulating DEGs, PRED also downregulated the expression of 16 NEGs in *mdx* muscles (compared to 7 for DMF treatment, Table 4). Notably, MMP13, which is crucial for muscle regeneration (43), was downregulated by 3-fold suggesting PRED may slow regenerative capacity. This may be a mechanism through which PRED constrains muscle size in DMD in addition to its atrophic effects mediated through antagonism of the insulin receptor (44). Collectively, our data suggest that DMF regulates a more specific ECM gene profile compared to PRED which could result in greater disease-modifying impact if these benefits were to persist longitudinally as DMD progresses.

**Table 4:** Effect of dimethyl fumarate (DMF) versus prednisone (PRED) treatment on normally expressed genes (NEGs) involved in extracellular matrix composition and cell adhesion in mdx compared to wild-type (WT) muscle (in order of most to least effected by PRED). Drug effect on NEG criteria is >1.5-fold regulation, *p*<0.05 from reference groups (*mdx* DMF and PRED v *mdx* VEH). Key: BMC: basement membrane constituents; CAM: cell adhesion molecule; CECMSC: collagens and extracellular matrix structural constituents; ECMM: extracellular matrix molecules; ECMP: extracellular matrix proteases; TR: transmembrane receptor. Blue shading indicates DMD disease seed genes.

| **NEG**<br>**(Mdx v WT)** |  | **DMF Effect** |  | **PRED Effect** |  | **Class** |
| --- | --- | --- | --- | --- | --- | --- |
| **Gene** | **Fold Reg.** | **Fold Reg.** | **p** | **Fold Reg.** | **p** |  |
| *MMP13* | 1.82 | -1.90 | 0.176 | ↓ **3.36** | ** | ECMM |
| *TNC* | 3.23 | ↓ **2.21** | * | ↓ **4.77** | **** | Other |
| *MMP3* | 1.94 | ↓ **2.23** | *** | ↓ 2.54 | 0.218 | ECMM |
| *SELP* | 1.39 | **-1.78** | * | ↓ **2.47** | * | CAM |
| *ITGB2* | 3.81 | -1.82 | 0.21 | ↓ 2.21 | 0.071 |  |
| *ITGAL* | 1.82 | -1.12 | 0.438 | ↓ **2.19** | **** |  |
| *SELE* | -1.43 | -1.43 | 0.132 | ↓ **2.18** | ** |  |
| *ADAMTS8* | -1.76 | 1.39 | 0.21 | **2.10** | * | ECMM |
| *ICAM1* | 2.03 | -1.33 | 0.061 | ↓ **2.08** | **** | CAM |
| *SELL* | -1.24 | -1.22 | 0.295 | ↓ 2.05 | 0.064 |  |
| *POSTN* | 2.82 | 1.17 | 0.46 | ↓ **2.01** | * | Other |
| *CD44* | 2.50 | -1.60 | 0.883 | **-1.85** | ** |  |
| *THBS1* | 1.74 | -1.25 | 0.407 | **-1.75** | ** |  |
| *ITGA3* | -1.50 | -1.29 | 0.103 | **-1.73** | * |  |
| *ENTPD1* | -1.01 | -1.11 | 0.449 | **-1.60** | ** |  |
| *COL6A1* | -1.14 | 1.16 | 0.229 | **-1.41** | * |  |
| *SGCE* | -1.55 | **1.27** | * | **-1.41** | * |  |
| *ITGA5* | 1.26 | **-1.35** | * | **-1.37** | * |  |
| *FN1* | 1.27 | 1.05 | 0.631 | **-1.32** | * |  |
| *CTGF* | 1.41 | **-1.80** | * | -1.05 | 0.685 |  |
| *COL5A1* | -1.17 | **1.20** | * | 1.11 | 0.533 |  |

We were also interested to understand whether DMF could temper muscle degeneration through Nrf2-mediated cytoprotection. Classical indicators of muscle histopathology were assessed in cryosectioned TA’s using haematoxylin and eosin (H&E) staining to derive the proportion of healthy (intact, peripherally nucleated fibres) and unhealthy (regenerating centronucleated fibres, degenerating fibres and inflammatory infiltrate) muscle (Fig 7). Healthy muscle was reduced, unhealthy muscle was increased and the unhealthy: healthy tissue ratio was higher in *mdx* compared to WT muscles (genotype effect; Fig 7A-C, E & G) consistent with dystrophin-deficient disease. DMF increased the overall proportion of healthy muscle Fig 7A) and reduced the overall proportion of unhealthy muscle in WT and *mdx* mice resulting in reduction of the unhealthy: healthy tissue ratio (Fig 7A-C, E-F & G-H). In contrast, PRED increased the proportion of unhealthy muscle (Fig 7B), specifically the relative area of centronucleated regenerating fibres compared to both *mdx* VEH and *mdx* DMF TA’s. (Fig 7D, G-I). Nuclear centration persists in regenerating/repairing myofibres for variable durations (up 94 weeks has been reported following chemotoxic muscle injury) and is associated with both satellite cell-mediated and myofibre autonomous repair mechanisms (45). Taken together with the 3-fold reduction in MMP13 expression (Table 3), our data suggest PRED slows muscle regeneration following damage, which may account for elevated NF-κB signalling (as observed in Fig 3). There was no evidence of global muscle atrophy due to genotype or treatment, although there was some redistribution of fibre size due to genotype (more smaller fibres in mdx groups) and PRED treatment (more medium sized fibres; see Supp Fig 4).

**Figure 7.**
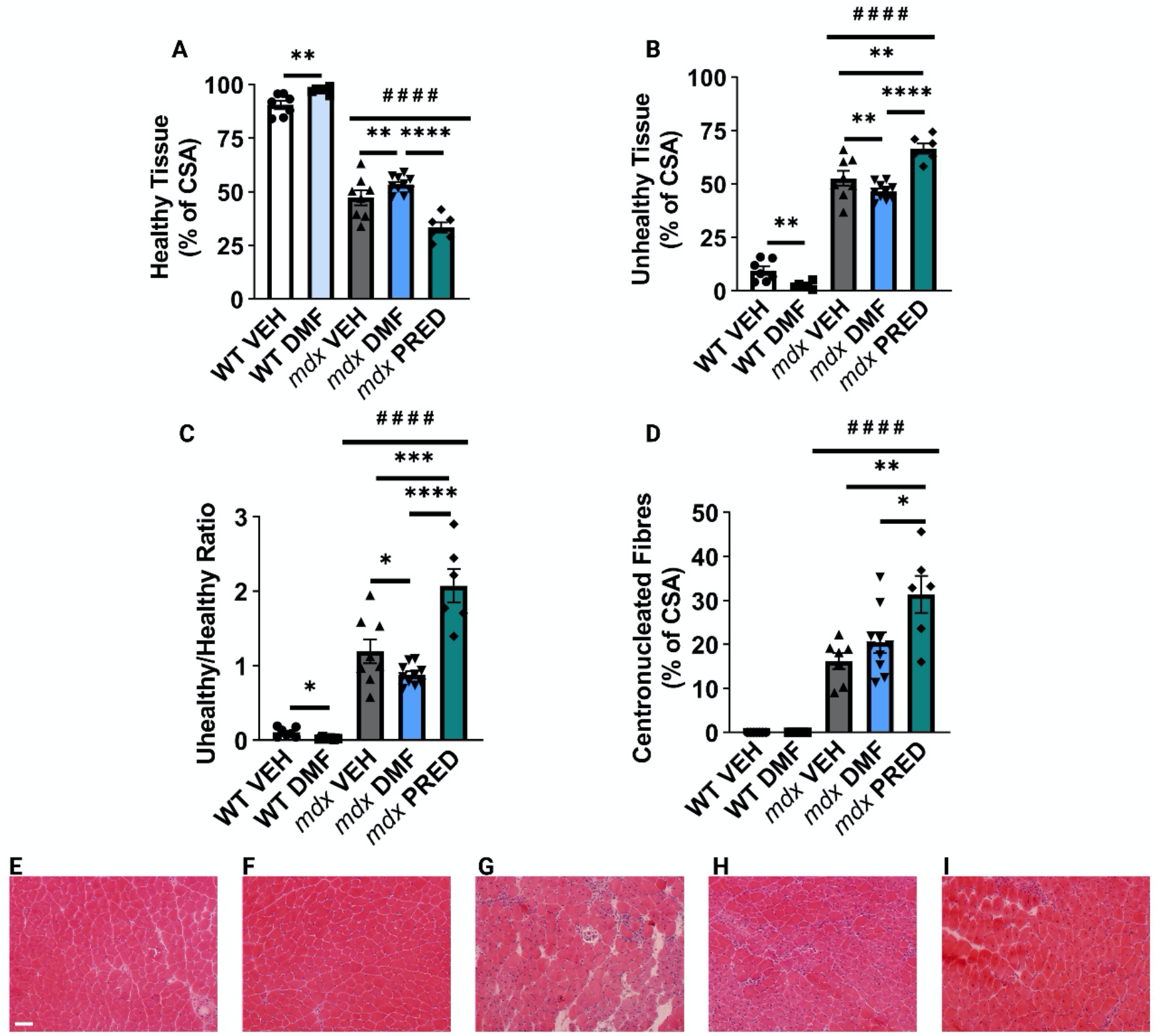
Dimethyl fumarate (DMF) improves *mdx* muscle histopathology. (A-I) Haematoxylin and eosin staining was utilised to assess the architecture of tibialis anterior (TA) muscles. (A) The proportion of healthy tissue was significantly reduced in *mdx* compared to wild-type vehicle (WT VEH) muscles (genotype effect) and was further reduced by prednisone (PRED) compared to *mdx* DMF treatment. DMF significantly increased the proportion of healthy tissue in WT and *mdx* TA compared to VEH. (B) The proportion of unhealthy tissue was increased in *mdx* muscles (group effect) but was reduced by DMF treatment yet exacerbated by PRED treatment. (C) The unhealthy: healthy tissue ratio was increased in *mdx* compared to WT muscles (group effect) and was reduced by DMF treatment. Conversely, PRED treatment increased the unhealthy: healthy tissue ratio compared to *mdx* VEH and DMF mice. (D) Centronucleated regenerating fibres were also increased in *mdx* muscles (group effect) and were increased by PRED treatment. Representative images of (E) WT VEH, (F) WT DMF, (G) MDX VEH, (H) MDX DMF and (I) MDX PRED TA muscles. Treatment effect: \**p*<0.05, \*\**p*<0.01, \*\*\**p*<0.001, \*\*\*\**p*<0.0001; genotype effect: ^####^*p*<0.0001

## Discussion

DMD is a difficult disease to clinically manage due to its pathological complexity. As a result, glucocorticoids have persisted as standard pharmacological care despite their low efficacy-to-side effect profile. The success of current and emerging gene-based therapies (e.g., exon skipping and gene transfer) aimed at restoring dystrophin expression, could ultimately depend on effectively treating the pathobiological milieu resulting from dystrophin deficiency, e.g., since ECM fibrosis might limit drug uptake into myofibres, and mitochondrial/endoplasmic reticular stress might constrain dystrophin protein synthesis at the ribosomal level. Experimental therapeutics targeting many different aspects of DMD pathobiology are in the clinical pipeline for these reasons (reviewed in (4)). However, thus far, there have been low translational success rates highlighting that a different approach to investigational drug selection is required.

In this study, re-purposed DMF was investigated for potential therapeutic benefit relative to standard care PRED due to its multi-modal effects (reviewed in (14)). We showed that both DMF and PRED improved muscle contractile function in EDL (composed predominantly of type II myofibres and the least damaged muscle as shown by EBD fluorescence), protected against sarcolemma damage (i.e., EBD uptake) of DIA muscles and modified an extensive list of inflammatory and ECM modulatory genes. However, only DMF consistently reduced sarcolemma damage and histopathology of hindlimb muscles and increase performance on clinically compatible function tests. The mechanisms involved are likely due to enhanced antioxidant defences and/or membrane stabilisation of basal lamina and ECM components. DMF increased protein levels of key anti-oxidative enzymes and normalised genes involved in basal lamina composition (e.g., *COL4α2*: ↓1.55-fold in *mdx* v WT, ↑ 1.28-fold by DMF *p*=0.065; Table 3) and the control of ECM environments (e.g., SPARC: ↓1.69-fold in *mdx* v WT, ↑ 1.27-fold by DMF *p*<0.05); Table 3) (46). In contrast, PRED conferred no protection on sarcolemma membranes of hindlimb muscles and appeared to slow regeneration during the acute severe disease phase in juvenile mice (46). Effective muscle regeneration depends on immune and transitional ECM signalling, and the potent pan immunosuppression conferred by PRED appears too strong to support expedient repair as recently indicated in juvenile *mdx* mice). Difficult to resolve in context of our histopathology data is that DMF could not abrogate clinically relevant haematologic biomarkers of DMD (e.g., CK or oxidised albumin) or improve force output of SOL despite conferring significant sarcolemma protection to this muscle. Rather both DMF and PRED drove systemic haematologic oxidation, which may be crucial to their immunomodulatory mechanisms of action. Smaller muscle fibres are remarkably resistant to damage in animal models and patients (e.g., the ocular muscles are well preserved during even advanced DMD) – DMF treatment made SOL muscles notably (although not statistically) smaller indicating protection against damage and degradation may come at the expense of force output. Longitudinal testing is required to determine whether DMF can alter systemic muscle wasting sufficient to modify disease course as indicated by plasma CK levels.

Disease and treatment omics data are useful tools to map pathobiologic pathways, identify disease drivers and potential drug targets, and enable efficient drug selection, especially for complex diseases like DMD (1). Recently, a 5 seed gene-driven interactome was identified through computational meta-analysis of studies quantitating muscle gene expression in DMD patients: drugs developed for MS, other autoimmune diseases and haematological cancers were revealed as ideal re-purposing candidates (28). DMF has proven efficacy in the clinical treatment of RRMS and the autoimmune disease, psoriasis. It is also currently being investigated for the treatment of acute myeloid leukemia (6). Our qRT-PCR gene array studies captured 4/5 of Lombardo *et al*.’s (28) disease module seed genes (see Fig 8), namely in order of hierarchy in the interactome: *MMP2*, *SPP1*, *TIMP1* and *FN1* (fibronectin 1). DMF effectively lowered the expression of all 3 fibrosis associated DEGs (*MMP2*, *SPP1* and *TIMP1*) but did not affect *FN1* (Figure 5, Table 3 and summarised in Fig 8). Conversely, PRED significantly downregulated *FN1*, whose upregulation is critical for augmenting muscle repair mechanisms (47). As a point of contrast, PRED better modulated expression of *TIMP1*, which is induced by cytokines and whose protein product inhibits MMP-mediated collagen degradation, while DMF was more effective against *SPP1*/OPN and *MMP*s (Fig 5G & Table 3). Although we didn’t assess *IGF-1* in this study, others have demonstrated (1) IGF-1 levels decrease as DMD progresses; and (2) DMF and analogous fumarate esters, increase *IGF-1* expression in neurons (48) while glucocorticoids notoriously reduce circulating IGF-1 levels (49). Collectively, these data suggest DMF could convey superior medicinal benefit than standard care corticosteroids to DMD patients as it does for MS patients by modulating a selective disease gene network over eliciting pan immunosuppression. Although corticosteroids are useful to abate the immune system during the initial phase of an MS relapse, only DMF can modulate the immune system over the long-term to reduce relapse rate (50). Longitudinal studies are required to decipher whether DMF or PRED will ultimately be more beneficial in modifying the natural history of DMD in *mdx* mice.

A distinct effect of DMF over PRED treatment was on mitochondrial function. Our data show acute DMF treatment augments mitochondrial respiration through increased substrate flux rather than through mitochondrial homeostasis signalling. Although we saw no evidence of altered mitochondrial function in juvenile *mdx* FDB fibres, mitochondrial anomalies are well reported in animal models of DMD and patients, including in muscle stem cells (reviewed in (34) and (51)). Complex I dysfunction has been reported in isolated mitochondria and fibres from *mdx* mouse muscle using different methodological approaches (36, 52, 53) and is probably linked to intracellular/mitochondrial calcium dysregulation as was reported mechanistically only recently (54). However, ATP production/phosphorylating respiration can be restored by re-routing respiration through Complex II (via addition of succinate and Complex I inhibitors (36, 52)). Our data suggest that DMF can achieve a similar mechanism during its end stage metabolism within the mitochondria TCA cycle, resulting in fatigue resistance to repetitive contraction in *mdx* muscles. Impaired mitochondrial homeostasis mechanisms, e.g., mitochondrial biogenesis and fission-fusion dynamics, are also described in dystrophin-deficient muscles and can be rescued via Nrf2 activation, although we saw no evidence of changes to crude protein markers with acute DMF treatment. Several mitochondria targeted therapeutics (e.g., ^(*+)*^epicatechin, MA-0211, elamipretide) are currently in clinical trials in DMD patients although none have shown efficacy in slowing disease course yet (55, 56). In fact, Santhera’s Phase IV idebenone trial was discontinued in 2020 due to futility (57) indicating that exclusively targeting mitochondria may be insufficient to slow the clinical course of DMD. DMF could represent a better alternative because it can modulate multiple drivers of DMD pathobiology including at the mitochondrial level.

Ultimately, the benefit of a drug to a patient population weighs efficacy against the unwanted side effect profile. PRED has an extensive side effect profile that restricts its therapeutic application to all patients, especially young patients and over the long term. In this study, we showed adverse effects on growth, fluid intake and spleen size in mice after only two weeks of PRED treatment. We commenced treatment prior to the onset of the acute severe MD phase at ~18d (i.e., treatment began at 14d age) to give maximum chance for therapeutic efficacy and attenuation of severe phasic DMD. Early treatment of DMD patients with glucocorticoids is recommended for the same reason (58). Ambulatory DMD patients treated with PRED show shorter stature, heavier weight and greater body mass index compared to steroid naïve patients and earlier commencement, higher dosage and longer duration are predictive of growth retardation (59). In contrast, DMF had no impact on growth or on the mass of any organ assessed in our mice. While it does have known side effects in humans – namely, flushing, gastrointestinal disturbances and in rare cases, leukopenia – we saw no adverse impact of DMF treatment on animal welfare parameters in this study e.g., on food consumption or susceptibility to pathogenic disease that would indicate gastrointestinal and/or leukopenia symptoms, respectively. Most of DMF’s side effects can be prevented or alleviated through dose ramping and more recently developed fumarate ester drugs, such as diroximel fumarate, have far fewer side effects (6). DMF was shown to be safe and efficacious in a 13 month multi-centre study in paediatric MS patients with no impact on growth (60) highlighting that if it were to impart PRED equivalent efficacy against DMD over the long term, it could prove a superior drug based upon side effect profile alone.

In summary, the data highlight acute DMF treatment as a robust modulator of the DMD disease module leading to extensive histopathological and functional benefits over PRED treatment. While DMF and standard care PRED both suppressed inflammatory genes, DMF selectively modulated disease-driving seed genes and had fewer apparent side effects in mice. Classical biomarker of DMD progression, plasma CK was not affected by either DMF or standard care PRED, highlighting that long term follow up pre-clinical studies are required to understand whether DMF can slow the progression of murine DMD better than PRED especially over the long term.

## Methods

### Animals

#### Breeding, Housing and Care

Dystrophin-positive C57BL/10ScSn WT mice and dystrophin-negative C57BL/10mdx (*mdx*) mice were bred from stock originally sourced from Animal Resources Centre (Western Australia, Australia) at the Western Centre for Health, Research and Education Animal Facility (Sunshine Hospital, Victoria, Australia), on a 12:12 hour light-dark cycle; 20-25°C; 40% humidity. Breeders and their litters were maintained in isolation separate to the main animal housing facility to limit stress. Animal welfare was monitored daily to accurately determine litter birth dates. Once born, litters remained in cages until weaning age (21 days). Thereafter, litters were housed in the main animal housing room in cages of 3-10 in treatment groups for the remainder of the study. From this point, food and water consumption and body weight was monitored daily. Dystrophin deficiency and dystrophin complexed protein downregulation in *mdx* mice was confirmed via western blot (Supp Figure 5).

#### Treatment Protocol

Homozygous littermates were randomly assigned to treatment groups at 14 days of age. WT and *mdx* mice were treated with either 0.5% methyl cellulose VEH (v/w) or ground DMF suspended in 0.5% methyl cellulose (v/w). A third cohort of *mdx* mice were treated with PRED suspended in 0.5% methyl cellulose (v/w). Animals were weighed daily (in the A.M.), and individual treatments were prepared relative to body weight to give a final daily dosage of either: 100mg.kg^−1^.day-1 DMF or 5 mg.kg^−1^.day^−1^ PRED. These dosages are consistent with previous pre-clinical studies of DMF for MS (6) and pre-clinical studies evaluating other drugs against *mdx* MD compared to PRED (61). Treatments were administered via oral gavage using a 21G gavage needle and animals were monitored for adverse events for ~5m post-gavage. Animals were treated up to and at 27 days of age (i.e., 14 days of treatment).

#### Functional Muscle Strength Testing

At 28 days, forelimb grip strength was measured using a commercial dynamometer (Bioseb, USA) over 3 consecutive efforts with 30 secs rest in between. The maximal effort (g) was used as absolute force (g) and was corrected for body mass (g/g). After 5 mins rest, mice were subjected to a four-limb hang test using a grid mesh system (custom) to assess whole body strength. Mice were excluded if they refused the test (hang <10 secs on 3 repeated attempts). The minimal holding impulse was calculated as body mass multiplied by absolute hang time.

#### Blood Biomarkers

After functional tests were performed on Day 28, tail ends were snipped, and blood was collected onto a PerkinElmer 226 Spot Saver RUO Card containing polyethylene glycol maleimide. Cards were stored with silica gel desiccant for transport to the University of Western Australia. Albumin was extracted into 0.05% tween 20 in 20mM phosphate with further binding to Cibacron Blue F3GA agarose, then eluted with 25μL of 1.4M NaCl in 20mM phosphate buffer pH 7.4. Gel electrophoresis, imaging and calculation of total albumin oxidation was performed as previously described by Lim *et al*. (62). On Day 28, mice were anaesthetised (4% induction, 2.5% maintenance isoflurane) and blood collected via terminal cardiac puncture into lithium heparin microtubes. Plasma was derived by centrifugation (3000G, 5 mins, 4°C) and CK was quantitated spectrophotometrically using a commercially available kit (Randox Laboratories, U.K.).

#### EBD Treatment

A separate cohort of (male and female) mice was utilised for EBD detection of skeletal muscle damage. This is because EBD interferes with standard histological staining protocols and fluorescence-based assays (such as extracellular flux) and may affect physiological parameters. Mice were injected with 1% EBD in saline (at 1% v/w) on Day 27, exactly 24 hours prior to tissue harvest and following the final gavage treatment.

#### Surgical Procedures

On Day 28, animals were weighed, deeply anaesthetised (4% induction and 2.5% maintenance isoflurane) and used for *ex vivo* experiments. Hind limb skeletal muscles were surgically excised in the following order: FDB, EDL, SOL, TA, plantaris, gastrocnemius and quadriceps. Organs/muscles were removed in the following order: DIA, heart, lungs, liver, spleen, and kidneys. Muscles and organs were weighed then processed for experiments.

### Metabolic Studies

#### Mitochondrial respiration & extracellular acidification

Isolated FDB fibres were prepared from whole FDB muscles as described by us previously, with modification (35). FDB muscles were incubated in pre-warmed dissociation media for 50 mins instead of 1.5 hours. Mitochondrial oxygen consumption and extracellular acidification rates were measured using a standard mitochondrial stress test on a Seahorse extracellular flux analyser (Agilent, USA).

#### CS activity

CS is the first enzyme of the Krebs cycle and an accepted biomarker of mitochondrial density. CS activity was determined as described by us previously (35).

### Ex vivo Muscle Contractile Function Studies

*Ex vivo* muscle contractile properties was performed as described by us previously on EDL and SOL using a Danish Myo Technology (DMT) system (Hinnerup, DNK (63).

### Muscle Histopathology

From EBD-treated mice, TA, EDL, SOL, and DIA strips were coated in OCT and snap frozen in chilled 2-methylbutane (in LN_2_; Sigma) and only TA was collected from all other non-EBD-treated mice. Processed muscles were serially cryo-sectioned (10 μm at −15 °C).

EBD sections were fixed in acetone (−15°C) and mounted with DPx (BDH, Poole, UK). Slides were imaged using TRITC filtered fluorescence microscopy at x40 magnification (BX53 Olympus Fluorescence Microscope). EBD positive fibres were quantitated using Image J (NIH, USA) and expressed relative to the muscle CSA.

TA cryosections from non-EBD treated mice were stained with a standard H&E protocol (35). To generate fibre size frequency distributions, all fibres on the cross section were individually traced on a Microsoft Surface tablet using ImageJ. For the quantitation of healthy versus non-healthy tissue, peripherally nucleated myofibres were distinguished from centronucleated myofibres and each group were counted and expressed relative to the total fibre number in the cross-section using Image J (64). Degenerating tissue was quantified as previously described (35).

SDH (Complex II) activity/capacity was also quantified in TA sections as described previously (35). Using Image J, images were deconvoluted (Haematoxylin and Periodic Acid of Schiff’s vector) and SDH activity positive intensity density quantified on the purple split relative to the total CSA.

Neutral lipid droplets were quantified in TA sections as described previously (35). Using Image J, images were deconvoluted (Fast Red: Fast Blue vector) and ORO positive intensity density quantified on the red split relative to the total CSA.

Unless otherwise stated, slides were imaged on a Zeiss Axio Imager Z2 scanning microscope at x200 magnification.

### Protein Expression

Western blotting was used to determine target engagement of Nrf2 by DMF and downstream cell signalling of the antioxidant and anti-inflammatory responses, mitochondrial dynamics, cell stress; as well as cytoskeletal proteins, in gastrocnemius as described by us previously (65). Primary antibodies used were: anti-Desmin (1:1000; #5332; CST), anti-DJ1 (1:1000; #5933; CST), anti-Dystrobrevin (1:500; #610766; BD Biosciences), anti-Dystrophin (1:500; ab15277; Abcam), anti-DRP-1 (1:1000; #8570; CST),) anti-HO-1 (1:1000; ADI-SPA-896; Enzo Life Sciences), anti-KEAP1 (1:1000; #8047; CST), anti-NF-κB (1:500; #8242; CST), anti-phospho-NF-κB (1:500; 3033; CST), anti-NQO1 (1:1000; #62262; CST), anti-Nrf2 (1:1000; #12721; CST), anti-OPA-1 (1:1000; #80471; CST), anti-phospho-p38 (1:750; #4511; CST), anti-p62 (1:1000; #5114; CST), anti-phospho-p62 (1:500; #95697; CST), anti-PGC-1α (1:1000; AB3242; Sigma-Aldrich), anti-SOD 1 (1:3000; ADI-SOD-101; Enzo Life Sciences), anti STAT3 (1:1000; #12640; CST), anti-phospho-STAT3 (1:2000; #9145; CST) and anti-TFAM (1:1000; ab252432; Abcam). Membranes were probed with a horseradish peroxidase-conjugated secondary antibody (1:5000; anti-rabbit IgG or 1:20,000; anti-mouse IgG, Vector Laboratories) in 5% not-fat milk powder in TBST (1h, RT) then stained with Coomassie Blue and normalised to total protein.

### Gene Arrays

Mature messenger RNA was isolated from quadriceps homogenates using the RNeasy1 mini kit (Qiagen, Hilden, Germany) according to manufacturer instructions. Cell lysates were transferred onto RNeasy mini-spin columns and DNA was removed using DNase digestion/treatment using RNase-Free DNase Set (Qiagen, Hilden, Germany.) The RNA Integrity Number (RIN) of all samples was quantified using an Agilent 2100 Bioanalyser and Agilent RNA 6000 nano kit (Agilent Technologies, Santa Clara, CA, USA). RIN values above 7.5 were used as the inclusion criterion for subsequent gene expression analysis. The concentration of RNA samples was measured using a Qubit RNA BR Assay (Invitrogen) in triplicate. Aliquots of each RNA sample were reverse transcribed to make complementary DNA (cDNA) using an RT2 first strand kit (Qiagen, Hilden, Germany) according to the manufacturer’s instructions. Quantitative real-time polymerase chain reaction (qRT-PCR) was performed using the Qiagen Mouse Inflammatory Response and Autoimmunity (PAMM-077Z) and Mouse Extracellular Matrix and Adhesion Molecules (PAMM-013Z) RT2 Profiler PCR arrays (Qiagen, Hilden, Germany) to evaluate relative gene/mRNA expression in WT compared to *mdx* and treated compared to untreated muscles. CT values were normalised based on a selection of reference genes (*ACTB*, *B2M*, *GAPDH*, *GUB*, H*SP90AB1*) and fold changes/regulation of gene expression were calculated using the 2^(−ΔΔCT) formula (GeneGlobe, QIAGEN). Differential expression (up and down regulation) genes were identified using the criteria of a >1.5-fold increase/decrease in gene expression, *p*<0.05 from reference group. Heatmaps were created using Log2 transformed Z scores.

### Statistics

Data are reported as mean ± SEM unless otherwise stated. Two-way ANOVA was used for all analyses with genotype and DMF treatment as factors. One way ANOVA was used to assess PRED treatment relative to other groups. Repeated measures analysis was used for body weight, food and water consumption, grip strength, muscle force frequency and fatigue studies. Tukey’s post hoc test was used for multiple comparisons. α was set at 0.05 and trends were reported at <0.1. DEGS criteria were >1.5-fold regulation, *p*<0.05. NEGS criteria were <1.5-fold regulation.

### Study Approval

Mice used in this study were generated from a breeding program approved by the Victoria University Animal Ethics Committee (AEETH 17-010, superseded by AEETH 20-005). At 14 days of age, mice were transferred to an approved experimental project (AEETH 17-007, superseded by 19-003). Animals were bred and cared for according to the Australian Code of Practice for the Care and Use of Animals for Scientific Purposes guidelines.

## Conflict of interest statement

ER has received consultancy fees from Santhera Pharmaceuticals and Epirium Bio outside the submitted work. NG has received grants and personal consultancy fees from Santhera Pharmaceuticals outside the submitted work. DF is a principal investigator for studies on spinal muscular atrophy sponsored by Hofmann-La Roche LTD. There are no other activities related to commercial companies. The authors declare they have no conflicting interests.

## Author Contributions

ER, CAT and SK drafted the manuscript. ER, CAT, DF and NG conceived the study. CAT, ER, SK, DAD, DGC, ND, and AB conducted experiments and analysis. NP, EJR, RB and LS conducted experiments. JT, PA, VA and JDH provided technical know-how and resources and interpreted data. PH provided intellectual input on the manuscript. All authors reviewed the manuscript.

## Acknowledgements

This work was supported by funding from Victoria University, University Children’s Hospital Basel, Muscular Dystrophy Association U.S.A. (Ideas grant: MDA871929), Duchenne UK and Save Our Sons. ER and CAT acknowledge outstanding support from Nicole Christie, Tricia Murphy and Dr Steven Holloway with animal care and breeding (VU Animal Services, WCHRE, Sunshine Hospital). ER and CAT thank Dr Craig Goodman (Centre for Muscle Research, University of Melbourne) for donation of the DJ-1 and DJ-1 Cys^106^ antibodies.

