## Supplemental Figures for "Dimethyl fumarate modulates the Duchenne muscular dystrophy disease program following short-term treatment in *mdx* mice"

### SUPPLEMENTAL MATERIAL

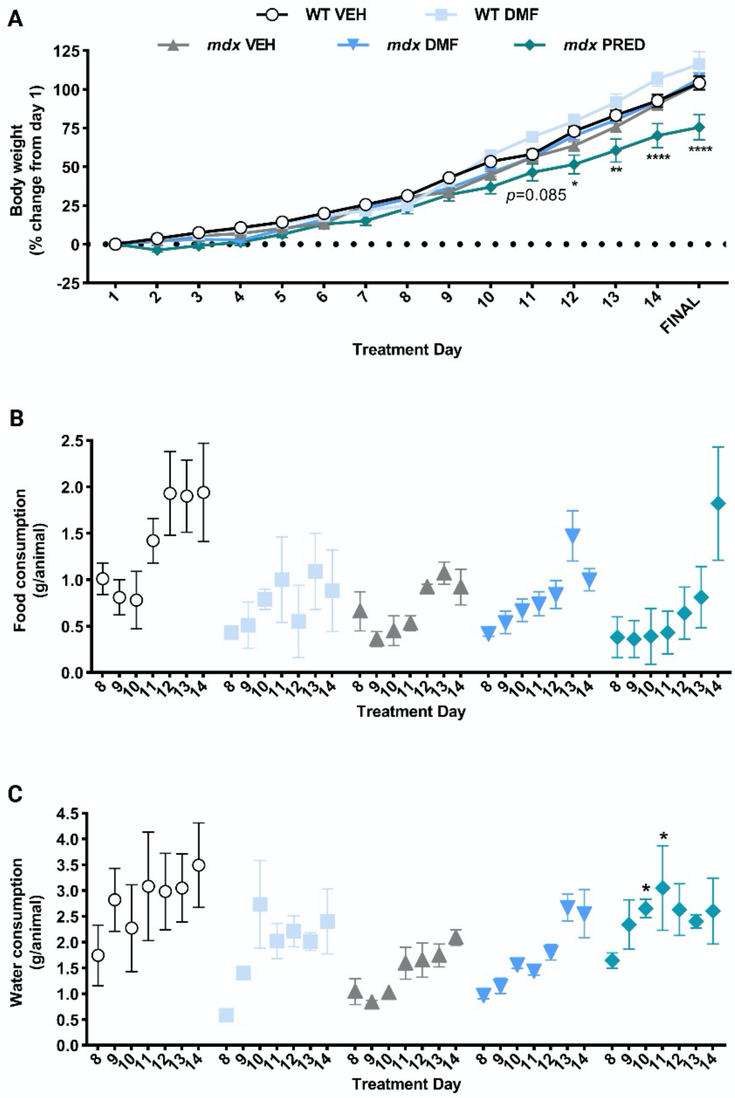


**Supplementary Figure 1: Daily dimethyl fumarate (DMF) treatment is safe in juvenile mice.** There was no effect of DMF treatment on growth (A), food (B) and water consumption (C) in wild-type (WT) or *mdx* mice. Prednisone (PRED) reduced growth rate of *mdx* mice from day 12 (trend from day 11) consistent with its known repressor effect on skeletal growth. Treatment effect: **p*<0.05, ***p*<0.01, *****p*<0.0001.

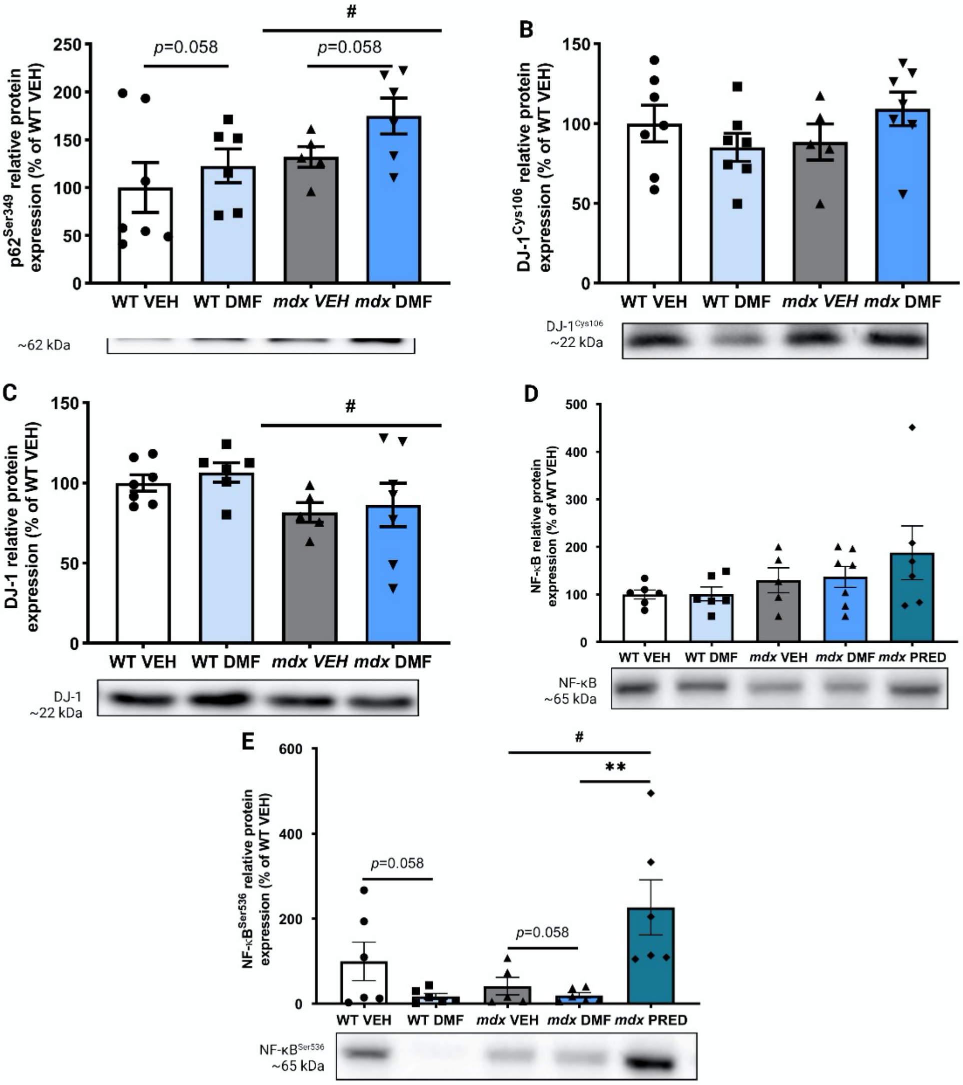


**Supplementary Figure 2: Protein expression of other proteins involved in Nrf2 activation and inflammation.** (A) Phosphorylated p62 (Serine 329) was elevated in *mdx* mice compared to wild-type (WT) mice with a trend for dimethyl fumarate (DMF) treatment to induce further phosphorylation observed. (B) While there was no difference in the phosphorylation of DJ-1 in any group, a reduction in total DJ-1 protein expression was detected in *mdx* mice (C). (D) Total NF-Kb protein was comparable in both WT and *mdx* mice (irrespective of treatment), however there was a trend for DMF treatment to reduce NF-Kb phosphorylation (Serine 536) in both WT and *mdx* muscles (E). Conversely, prednisone (PRED) treatment increased nf-κB phosphorylation compared to *mdx* vehicle (VEH) and DMF treated mice. Treatment effect: **$p<0.01$; genotype effect: $^{\#}p<0.05$.

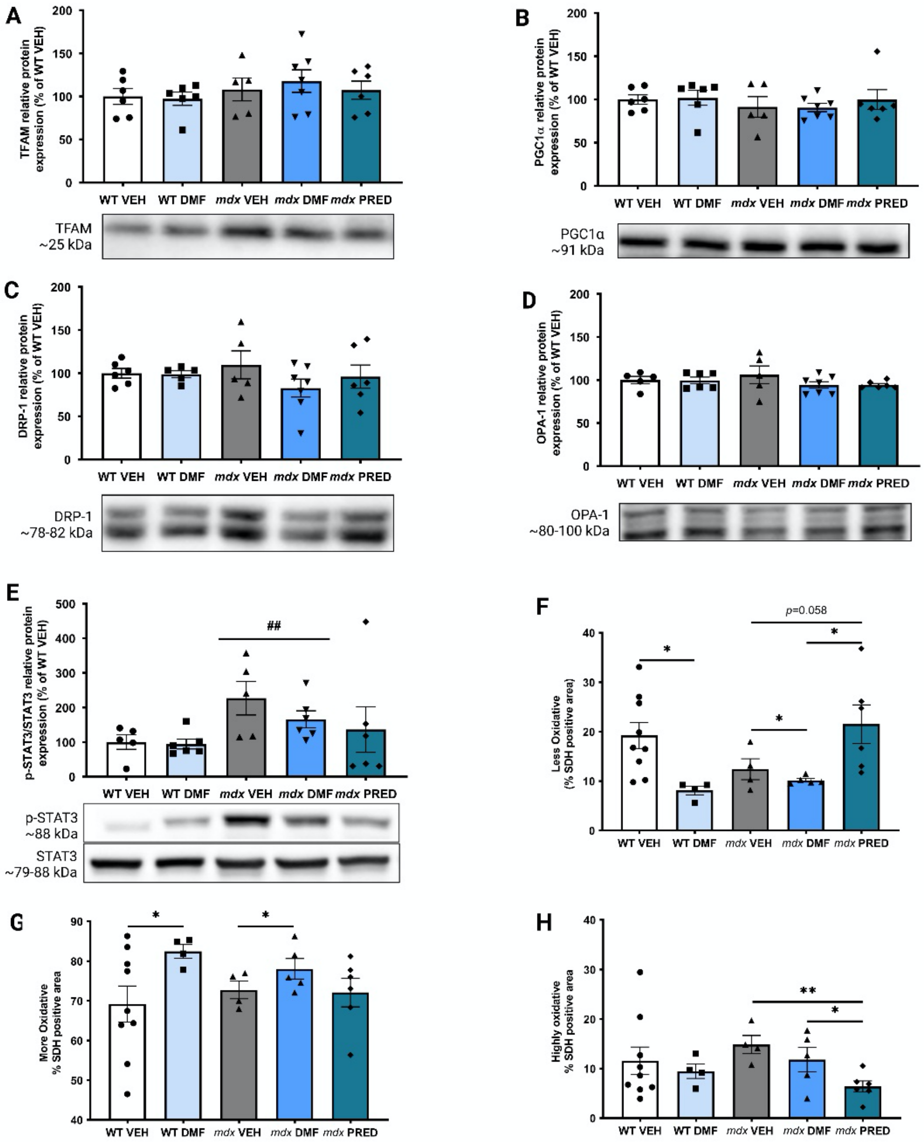


**Supplementary Figure 3: Dimethyl fumarate (DMF) does not alter mitochondrial protein expression but does augment the oxidative capacity of *mdx* muscles.** No difference in TFAM (A), PGC1α (B), DRP-1 (C) or OPA-1 (D) expression was observed in either wild-type (WT) or *mdx* mice, irrespective of treatment (vehicle (VEH), DMF or prednisone (PRED)). (E) The ratio of phosphorylated to total STAT3 protein expression was higher in *mdx* mice compared to WT muscle. DMF treatment reduced the area of less oxidative muscle fibres (F) and increased the area of more oxidative fibres in both WT and *mdx* mice (G). PRED treatment reduced the area of highly oxidative muscle compared to *mdx* VEH and DMF mice (H). Treatment effect: *$p<0.05$, **$p<0.01$; genotype effect: $^{\#\#}p<0.01$.

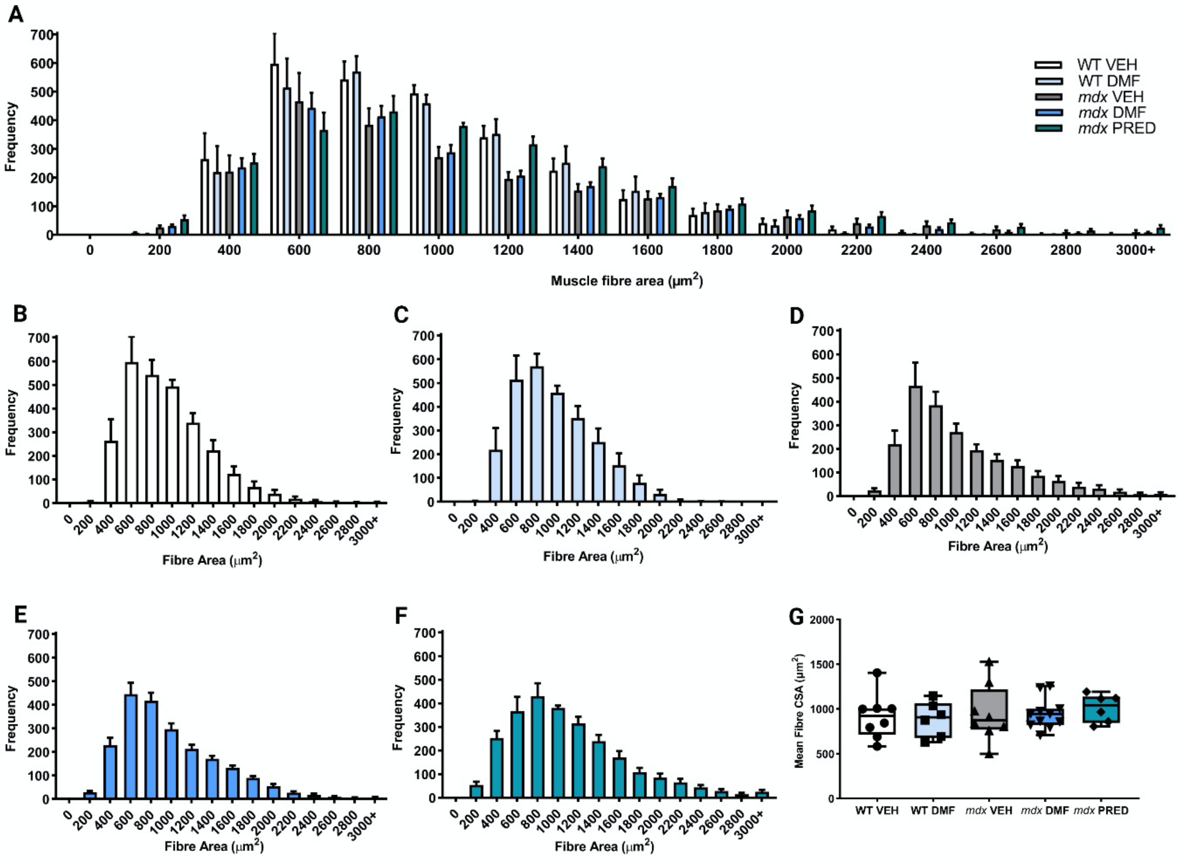


**H**

| Bin | 200-399 | 600-799 | 800-999 | 1000-1199 | 1200-1399 | 1400-1599 | 2400-2599 | 2600-2799 |
|---|---|---|---|---|---|---|---|---|
| **WT v *mdx* (Group Effect)** | # | 0.08 | # | #### | #### | ### | 0.06 | 0.095 |
| **WT VEH v WT DMF** | | | | | | | | |
| ***mdx* VEH v *mdx* DMF** | | | | | | | | |
| ***mdx* VEH v *mdx* PRED** | | * | | 0.09 | * | | | |
| ***mdx* DMF v *mdx* PRED** | | 0.09 | | | * | | | |

**Supplementary Figure 4: Dimethyl fumarate (DMF) does not impact the cross-sectional area (CSA) of tibialis anterior muscle fibres.** (A) A comparison of the fibre size distribution of wild-type (WT) vehicle (VEH), WT DMF, *mdx* VEH, *mdx* DMF and *mdx* prednisone (PRED) tibialis anterior with individual group fibre size histograms (B-F). (G) The mean CSA of tibialis anterior fibres was comparable across the groups, despite differences in the number of fibres sized between 200-399, 600-799 (trend), 800-999, 1000-1199, 1200-1399, 1400-1599, 2400-2599 (trend) and 2600-2799 (trend) detected between *mdx* and WT mice (H). PRED, but not DMF, altered the distribution of fibres in *mdx* mice. Treatment effect: *$p$<0.05; genotype effect: $^{\#}p$<0.05, $^{\#\#\#}p$<0.001, $^{\#\#\#\#}p$<0.0001.

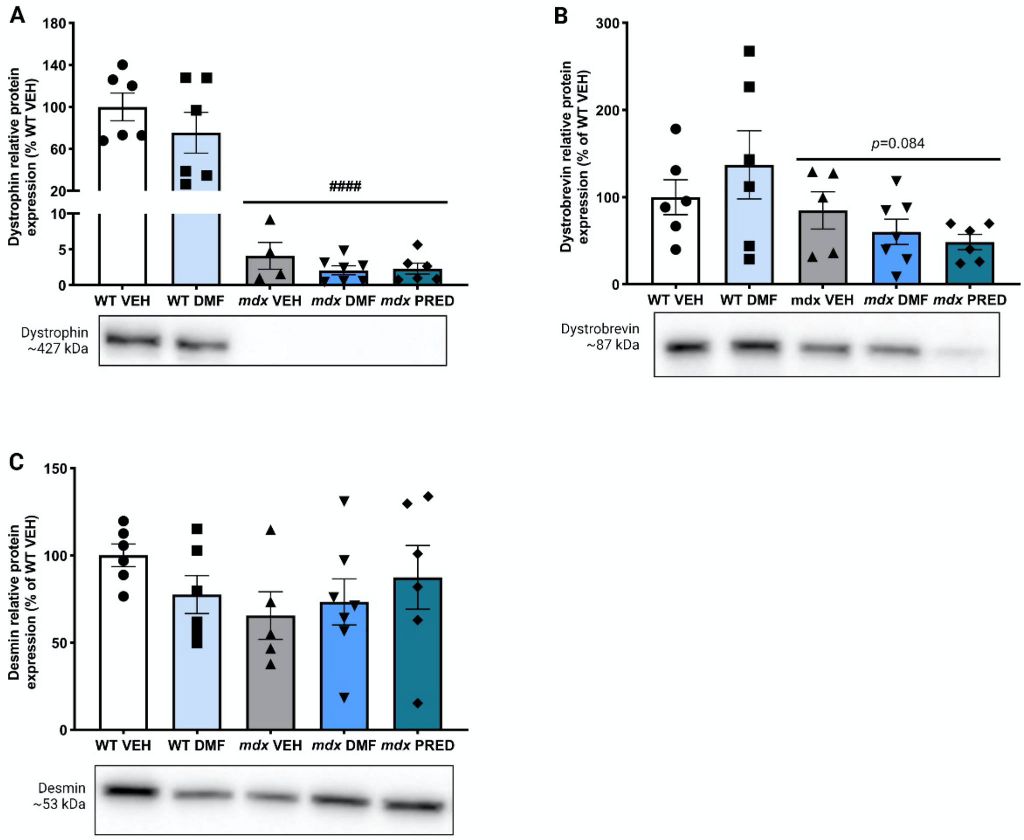


**Supplementary Figure 5: Dimethyl fumarate (DMF) does not alter the expression of the cytoskeletal proteins dystrophin, dystrobrevin or desmin.** (A) As expected, dystrophin expression was ablated in *mdx* muscle (compared to wild-type (WT)). Dystrophin-associated glycoprotein complex protein, dystrobrevin also trended toward reduction in *mdx* muscle (B). (C) Desmin expression was comparable between groups. Genotype effect: $^{\#\#\#\#}p<0.0001$.
